# Dimer-monomer transition defines a hyper-thermostable peptidoglycan hydrolase mined from bacterial proteome by lysin-derived antimicrobial peptide-primed screening

**DOI:** 10.1101/2024.04.26.591248

**Authors:** Li Zhang, Fen Hu, Zirong Zhao, Xinfeng Li, Mingyue Zhong, Jiajun He, Fangfang Yao, Xiaomei Zhang, Yuxuan Mao, Hongping Wei, Jin He, Hang Yang

## Abstract

Phage-derived peptidoglycan hydrolases (i.e., lysins) are considered promising alternatives to conventional antibiotics due to their direct peptidoglycan degradation activity and low risk of resistance development. The discovery of these enzymes is often hampered by the limited availability of phage genomes. Herein, we report a new strategy to mine active peptidoglycan hydrolases from bacterial proteomes by lysin-derived antimicrobial peptide-primed screening. As a proof-of-concept, five <u>p</u>eptidoglycan <u>h</u>ydrolases from the *<u>A</u>cinetobacter <u>b</u>aumannii* proteome (PHAb7-PHAb11) were identified using PlyF307 lysin-derived peptide as a template. Among them, PHAb10 and PHAb11 showed potent bactericidal activity against multiple pathogens even after treatment at 100°C for 1 hour, while the other three were thermosensitive. We solved the crystal structures of PHAb8, PHAb10, and PHAb11 and unveiled that hyper-thermostable PHAb10 underwent a unique folding-refolding thermodynamic scheme mediated by a dimer-monomer transition, while thermosensitive PHAb8 formed a monomer. Two mouse models of bacterial infection further demonstrated the safety and efficacy of PHAb10. In conclusion, our antimicrobial peptide-primed strategy provides new clues for the discovery of promising antimicrobial drugs.

## Introduction

Despite decades of concerted action from all over the world, antimicrobial resistance remains a growing global threat today (Nathan 2020). The misuse and overuse of antimicrobials leads to their accumulation in the environment, eventually increasing the chances of bacterial pathogens to acquire resistance (Van Boeckel, Pires et al. 2019, Rawson, Ming et al. 2020). Therefore, the concept of “One Health” has been proposed to address the alarming levels of antimicrobial resistance (McEwen and Collignon 2018, Hernando-Amado, Coque et al. 2019). In recent decades, the number of bacteria resistant to first-line drugs, as well as second and third line drugs, has been increasing. Some of these bacteria have even been identified as multi-drug resistant, extensively drug resistant, or pan-drug resistant isolates (Talaat, Zayed et al. 2022). For example, methicillin-resistant *Staphylococcus aureus*, vancomycin-resistant *S. aureus*, vancomycin-resistant enterococcus, colistin-resistant *Enterobacteriaceae*, third-generation cephalosporin-resistant *Escherichia coli* and *Klebsiella pneumoniae*, carbapenem-resistant *E. coli*, *K. pneumoniae*, and *Pseudomonas aeruginosa*, and multidrug-resistant *Acinetobacter baumannii* pose a serious threat and are the leading cause of morbidity and mortality in human infectious diseases (Brink 2019). In addition, the global medical need for new antimicrobials has not yet been fully addressed.

Considerable evidence shows that bacterial peptidoglycan is a promising target for the development of antimicrobial agents with a low risk of resistance development. A well-documented example in this regard are the peptidoglycan hydrolases, also known as lysins, derived from phages. In recent years, lysins have been shown to be efficient alternatives to traditional antibiotics for infections caused by Gram-positive bacteria in various animal models (De Maesschalck, Gutierrez et al. 2020, Schmelcher and Loessner 2021). Encouragingly, several lysins targeting *S. aureus* have been evaluated in clinical trials (Theuretzbacher and Piddock 2019). However, the development of lysins targeting Gram-negative bacteria is relatively recent, mainly because the bacterial outer membrane prevents lysins from accessing their peptidoglycan substrates. In the last decade, several strategies have been developed to enable lysin to overcome the outer membrane barrier and directly target Gram-negative bacteria. For instance, Artilysins are constructed by fusion of cationic nonapeptides to effectively eradicate resistant and persistent *A. baumannii* (Defraine, Schuermans et al. 2016); Innolysins are engineered by fusion of receptor binding proteins to displays bactericidal activity against *E. coli* that was resistant to third-generation cephalosporins (Zampara, Sorensen et al. 2020); bioengineered lysin-bacteriocin fusion molecule, i.e., Lysocins, are able to deliver lysin across the outer membrane of Gram-negative bacteria and show selective anti-Pseudomonal activity (Heselpoth, Euler et al. 2019). In addition, the establishment of high-throughput mining or engineering strategies has accelerated the discovery of active lysins against Gram-negative pathogens, e.g., the VersaTile-driven platform, which can rapidly screen engineered lysins active against *A. baumannii* from tens of thousands of combinations by integrating new DNA assembly methods and iterative screening procedures (Gerstmans, Grimon et al. 2020). However, compared with the current progress in the clinical translation of lysins against Gram-positive bacteria, the discovery of lysins against Gram-negative bacteria that meet the needs described in the WHO priority pathogen list is still urgently needed (Briers and Lavigne 2015, Lai, Chen et al. 2020).

Additionally, the discovery of phage-derived peptidoglycan hydrolases has also been hampered by limited sources of published phage genome data. Although not yet tested extensively, recent progress in lysins targeting Gram-negative bacteria have demonstrated a clue linking antibacterial activity to their internal antimicrobial peptides (Vazquez, Garcia et al. 2021), by which lysins cross the bacterial outer membrane and contact its underlying peptidoglycan substrates (Lood, Winer et al. 2015, Thandar, Lood et al. 2016, Li, Jiang et al. 2021). These observations prompted us to consider whether antibacterial peptidoglycan hydrolases or their homologues could be mined from bacterial proteomes using lysin-derived antimicrobial peptide as template. As a proof-of-concept, we here used the antimicrobial peptide P307 derived from PlyF307 lysin (Lood, Winer et al. 2015) as a template to search for <u>p</u>eptidoglycan <u>h</u>ydrolases from the *<u>A</u>. <u>b</u>aumannii* proteome database (PHAbs, Fig. S1). Two of them (PHAb10 and PHAb11) were found to be hyperthermally stable even after treatment at 100°C for 1 hour, which is mediated by a unique monomer-dimer swapping. Furthermore, both were active against a wide variety of clinically relevant bacteria *in vitro* and in mouse infection models. This observation supports our hypothesis that an antimicrobial peptide-primed mining strategy is feasible for discovering new peptidoglycan hydrolases with high bactericidal activity and desirable physicochemical properties from bacterial proteomes.

## Results

### Screening of putative peptidoglycan hydrolases from the *A. baumannii* proteome based on a lysin-derived antimicrobial peptide-primed mining strategy

Since a variety of lysins active against Gram-negative bacteria are shown to harbor an N-or C-terminal antimicrobial peptide, this led us to speculate that antimicrobial peptide could be used as template to mine new peptidoglycan hydrolases. To prove this hypothesis, P307, a well-documented antimicrobial peptide against *A. baumannii* from PlyF307 lysin (Lood, Winer et al. 2015, Thandar, Lood et al. 2016), was used as the template to mine putative <u>p</u>eptidoglycan <u>h</u>ydrolases from *<u>A</u>. <u>b</u>aumannii* (PHAb) proteome databases publicly available in NCBI (Fig. 1a). Finally, 204 hits were identified and further divided into 5 clades based on phylogenetic analysis. Clade I contained 162 hits with high similarity (> 85%) to the AcLys lysin from *A. baumannii* 5057UW prophage (Fig. 1b), and clade II-V contained 42 hits with relatively lower homology (40-77%) to each other (Fig. 1b and Fig. S2a).

**Fig. 1.**
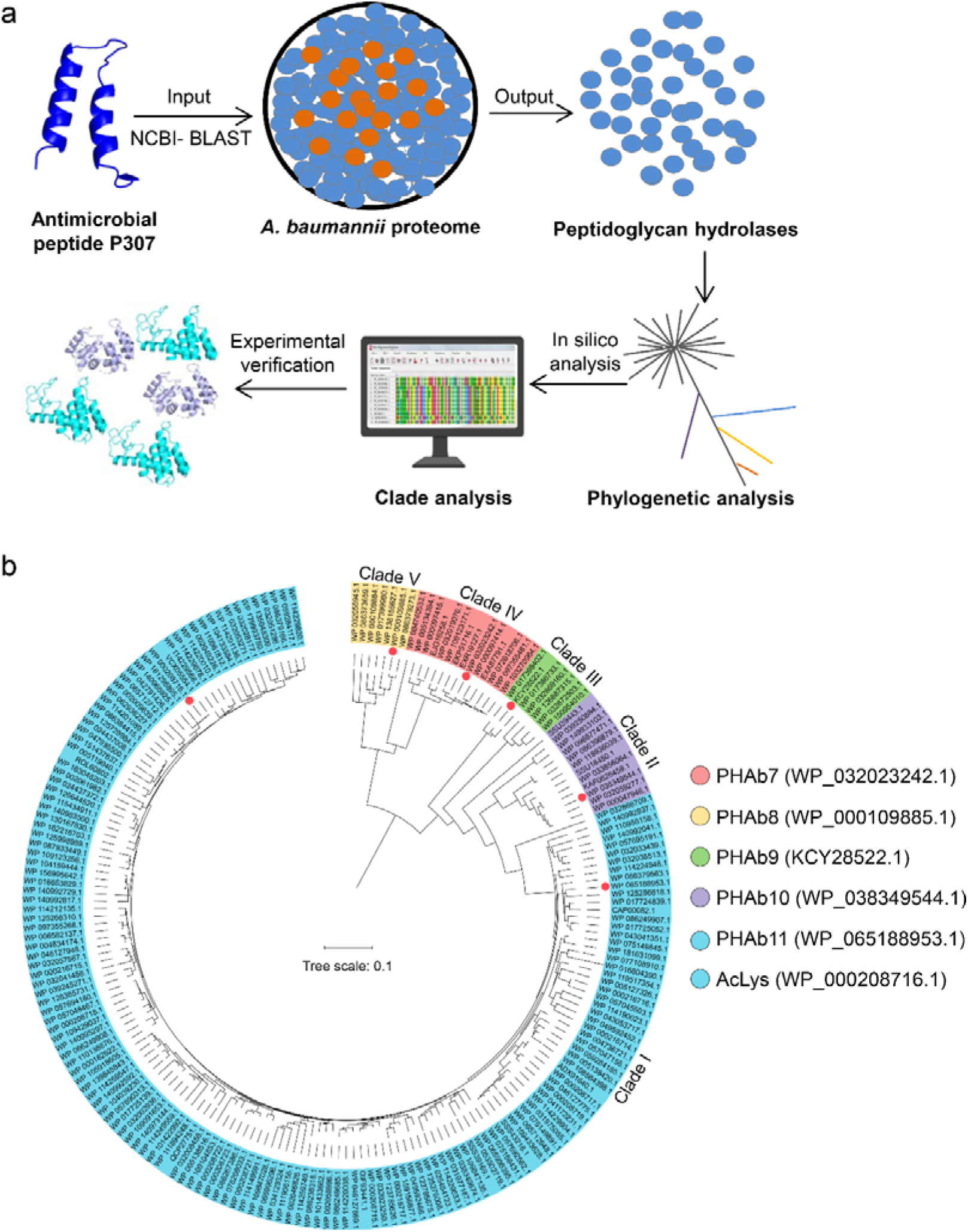
Screening of putative peptidoglycan hydrolases from the *A. baumannii* proteome. (a) Workflow for the development of an integrated system for peptidoglycan hydrolase screening. (b) Phylogenetic analysis of putative peptidoglycan hydrolases from the *A. baumannii* proteome database. Multiple sequence alignments are performed using MEGAX software with ClustalW algorithm, and the Neighbor-joining method is used to construct a phylogenetic tree. Classified clades are labeled in different colors and representative peptidoglycan hydrolases from each clade are pointed by colored circles.

Due to the high similarity of amino acid sequences within each clade, we selected a putative peptidoglycan hydrolase from each clade for further characterization by analyzing the selectivity priority in evolution and the overall physicochemical properties (water solubility, hydrophilicity, hydrophobicity, charge, flexibility, and rigidity) of each sequence *in silico*. Finally, WP_065188953.1 from clade I, WP_0383495441.1 from clade II, KCY28522.1 from clade III, WP_000109885.1 from clade IV, and WP_032023242.1 from clade V were shortlisted and renamed as PHAb11, PHAb10, PHAb9, PHAb8, and PHAb7, respectively, for further investigation (Fig. 1b). Further homology analysis revealed that these five peptidoglycan hydrolases showed great diversity with the structured AcLys lysin and the original PlyF307 lysin (Fig. S2b).

### PHAb10 and PHAb11 are highly thermostable peptidoglycan hydrolases with a broad-spectrum of action

To test the activity of these PHAbs, their coding sequences were chemically synthesized, cloned into a vector, and expressed in *E. coli* cells. As expected, all five PHAbs were well expressed as soluble proteins in *E. coli* BL21(DE3) with high purity (Fig. S3a). Next, we examined the hydrolytic activity of these 5 PHAbs on peptidoglycan derived from *A. baumannii* 3437 by a halo assay (Vander Elst, Linden et al. 2020). The results showed that all PHAbs form clear zones on the autoclaved *A. baumannii* lawns even at a low concentration of 2.5 μg/ml (Fig. S3b), indicating that these PHAbs are active against peptidoglycan of *A. baumannii*. To further confirm the ability of these PHAbs against peptidoglycan from different bacteria, we measured the area of clear zones generated by different concentrations of PHAbs on each lawn of two isolates of *A. baumannii*, *P. aeruginosa*, and *E. coli*. Encouragingly, these PHAbs were active against peptidoglycans from all strains tested and formed a clear halo (Fig. 2a). Among these PHAbs, PHAb10 and PHAb11 showed larger size of clear zones under each concentration tested (Fig. 2a), indicating that these two PHAbs might have stronger peptidoglycan hydrolyzing activity. To prove this, we further evaluated the bactericidal activity of these PHAbs against each isolate of *A. baumannii*, *P. aeruginosa*, and *E. coli* by a log-killing assay. Consistent with the halo assay, PHAb10 and PHAb11 exhibited outstanding bactericidal activity against all strains tested, with a 3.5-4.6 log reduction in viable bacterial counts after treatment for 1 hour with 50 μg/ml of each enzyme (Fig. 2b). Like other lysins, the potent bactericidal activity of PHAb10 and PHAb11 was time-dependent. After treatment with 50 μg/ml of either enzyme, a [0.9 log reduction in viable *A. baumannii* 3437 for both enzymes were observed within the first 30 seconds, with a substantial reduction to 3.1 logs (from 6.0 log to 2.9 log) for PHAb10 and 2.5 logs (from 6.0 log to 3.5 log) for PHAb11 after 15 minutes (Fig. 2c).

**Fig. 2.**
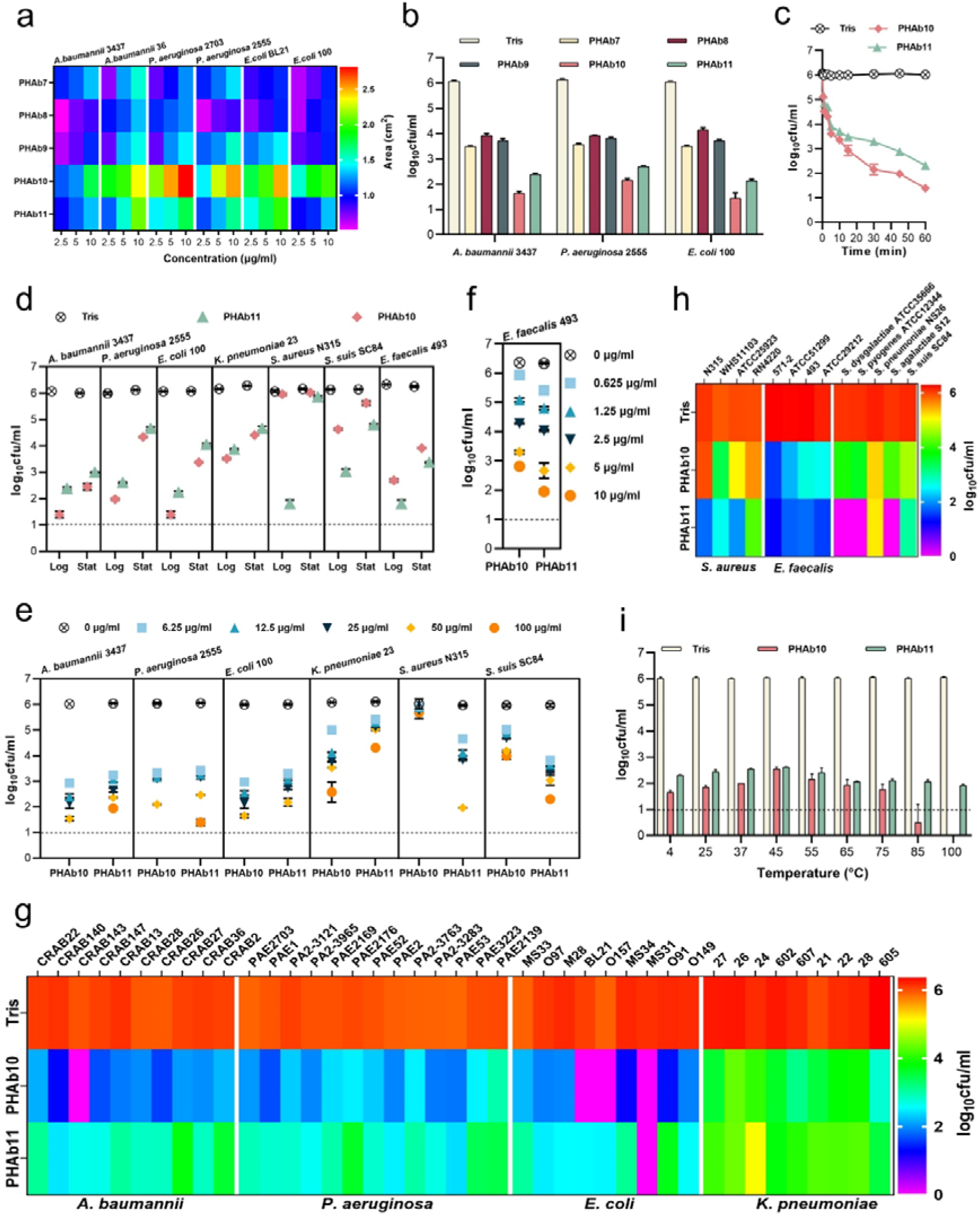
PHAb10 and PHAb11 are highly thermostable peptidoglycan hydrolases with a broad-spectrum of action. (a) Heat map showing the halo size formed by different concentrations of peptidoglycan hydrolases on different bacterial lawns. (b) Bactericidal activity of five peptidoglycan hydrolases (50 μg/ml) in 20 mM Tris-HCl (pH 7.4) at 37°C for 1 hour against different exponential bacteria. (c) Time-dependent bactericidal activity of 50 μg/ml PHAb10 or PHAb11 against exponential *A. baumannii* in 20 mM Tris-HCl (pH 7.4). (d) Bactericidal activity of 50 μg/ml PHAb10 or PHAb11 in 20 mM Tris-HCl (pH 7.4) at 37°C for 1 hour against different bacteria in exponential and stationary phases. (e-f) Dose-dependent bactericidal activity of PHAb10 and PHAb11 against multiple exponential bacteria in 20 mM Tris-HCl (pH 7.4) at 37°C for 1 hour. (g-h) Susceptibility of various Gram-negative (g) and Gram-positive (h) bacterial strains to PHAb10 and PHAb11. Exponential cultures of each bacterium are treated with 50 μg/ml PHAb10 or PHAb11 for 1 hour at 37°C and residual viable bacterial cells are counted by plating serial dilutions onto agar plates. For *E. faecalis*, 10 μg/ml of each peptidoglycan hydrolase is used. (i) Thermal stability of PHAb10 and PHAb11. Each enzyme is stored at different temperatures for 1 hour, cooled to room temperature, and then incubated with exponential *A. baumannii* 3437 in 20 mM Tris-HCl (pH 7.4) at a final concentration of 50 μg/ml at 37°C for 1 hour. Viable bacteria are counted after each treatment by plating serial dilutions on LB agar. Dash lines represent the limit of detection and data below the limit of detection is not shown.

Since previous reports documented the diverse performance of lysins against Gram-negative and Gram-positive bacterial species in exponential and stationary phases (Lood, Winer et al. 2015), we therefore examined the bactericidal activity of PHAb10 and PHAb11 against different bacteria in different growth phases. Interestingly, robust bactericidal activity was observed for both enzymes in exponential and stationary phase cultures of *A. baumannii*, *P. aeruginosa*, *E. coli*, *K. pneumoniae*, *S. suis*, and *E. faecalis* (Fig. 2d). It should also be noted that exponential bacteria were more susceptible to these two enzymes than stationary bacteria, similar to observations reported elsewhere (Lood, Winer et al. 2015). However, for *S. aureus*, detectable susceptibility to PHAb11 was only observed in its exponential phase (Fig. 2d). In addition, PHAb10 and PHAb11 exhibited dose-dependent killing against sensitive Gram-negative and Gram-positive strains tested. Specifically, treatment with 6.25 μg/ml PHAb10 for 1 hour resulted in a 3 log reduction in *A. baumannii*, a 2.7 log reduction in *P. aeruginosa*, a 3 log reduction in *E. coli*, 1.08 log reduction in *K. pneumoniae*, and a 0.94 log reduction in *S. suis*, and treatment with 6.25 μg/ml PHAb11 for 1 hour caused a 2.8 log reduction in *A. baumannii*, a 2.6 log reduction in *P. aeruginosa*, a 2.7 log reduction in *E. coli*, a 0.69 log reduction in *K. pneumoniae*, a 1.3 log reduction in *S. aureus*, and a 2.15 log reduction in *S. suis* (Fig. 2e). Interestingly, *E. faecalis* was more sensitive to PHAb10 and PHAb11, showing a 3.8 log and a 4.6 log reduction respectively after treatment with 10 μg/ml of each enzyme for 1 hour (Fig. 2f).

To further confirm the bactericidal activity of PHAb10 and PHAb11, the susceptibility of various clinical isolates such as *A. baumannii*, *P. aeruginosa*, *E. coli*, *K. pneumonia*, *S. aureus*, and *E. faecalis,* as well as representative isolates of *S. dysgalactiae, S. agalactiae*, *S. pyogenes*, *S. pneumoniae*, and *S. suis* to both enzymes were examined by log-killing assay. Our results showed that all tested strains, whether antibiotic-resistant or not, were sensitive to PHAb10 and PHAb11 (Fig. 2g-h). Importantly, Gram-negative bacteria were more sensitive to PHAb10, with 3.9-6.0 log reductions for *A. baumannii* isolates, 3.7-5.0 log reductions for *E. coli* isolates, 2.4-4.3 log reductions for *P. aeruginosa* isolates, and 1.1-3.6 log reductions for *K. pneumoniae* isolates (Fig. 2g). In contrast, PHAb11 was more bactericidal than PHAb10 against Gram-positive bacteria, causing 1.8-4.3 log reductions for *S. aureus* isolates, 4.5-5.0 log reductions for *E. faecalis* isolates, and 1.0-6.1 log reductions for various streptococcal strains (Fig. 2h).

Further biochemical characterization showed that PHAb10 and PHAb11 remained highly active at pH 5.0-10.0, and the maximum bactericidal activity was observed at pH 7.0 for both enzymes (Fig. S3c). Similar to other previously reported lysins (Yu, Frank et al. 2018, Kim, Jin et al. 2020), PHAb10 and PHAb11 were sensitive to NaCl and urea in a dose-dependent inhibitory manner (Fig. S3d). Surprisingly, both enzymes retained outstanding bactericidal activity after heat treatment at temperature up to 100°C for 1 hour (Fig. 2i). Indeed, after 1 hour treatment at 100°C, the bactericidal activity was improved slightly for both enzymes. In contrast, PHAb7, PHAb8, and PHAb9 completely lost their activity after treatment at 70°C for 1 hour (Fig. S3e). Together, these observations showed that PHAb10 and PHAb11 were highly thermostable peptidoglycan hydrolases with a broad spectrum of action.

### Different action mechanisms of PHAb10 and PHAb11 in killing Gram-negative and Gram-positive bacteria

Due to the natural barrier of the outer membrane, several peptidoglycan hydrolases (i.e., lysins) have been reported to have no or low bactericidal activity against Gram-negative bacteria in the absence of outer membrane penetrating agents (Briers and Lavigne 2015, Ghose and Euler 2020, Lai, Chen et al. 2020). However, for most of these lysins, their activity are almost exclusively restricted to Gram-negative bacteria (Guo, Feng et al. 2017, Khan, Gondil et al. 2021), raising questions about the mechanisms of action of PHAb10 and PHAb11 in killing both Gram-negative and Gram-positive bacteria. Structural prediction analysis showed that PHAb10 contains a putative lysozyme catalytic domain (amino acids 1-110, P10-Lys) and a C-terminal cationic peptide (amino acids 111-149, P10-CP), whereas PHAb11 contains an N-terminal cationic peptide with unknown function (amino acids 1-36, P11-NP), followed by a putative lysozyme catalytic domain (amino acids 37-145, P11-Lys, with 68.5% sequence similarity with P10-Lys) and a C-terminal cationic peptide (amino acids 146-184, P11-CP, which shows 94.9% sequence similarity with P10-CP) (Fig. 3a). Studies have shown that the cationic region of phage lysins can facilitate their penetration into the outer membrane, thereby fulfilling their bactericidal activity against Gram-negative bacteria (Lood, Winer et al. 2015, Yu, Frank et al. 2018, Li, Jiang et al. 2021). To examine whether it is the reason for the killing ability of PHAb10 and PHAb11 against Gram-negative bacteria, we further analyzed *in silico* the physicochemical properties of these putative cationic peptides harbored by these two enzymes and found that all three peptides, that is, P10-CP, P11-CP, and P11-NP might act as antimicrobial peptides (Table S1). Therefore, we synthesized these peptides and examined their antibacterial activity by log-killing assay. As shown in Fig. 3b, we observed bactericidal activity in P10-CP and P11-CP against *A. baumannii*, *P. aeruginosa*, *S. aureus*, and *E. faecalis* in both the exponential and stationary phases. However, rare lytic activity was observed in P11-NP, suggesting that its function remains to be established. Notably, P11-CP was more robust than P10-CP (Fig. 3b), which might be due to the presence of an additional positively charged amino acid (i.e., lysine) in P11-CP (Table S1). More importantly, the putative lysozyme catalytic domains of two enzymes, P10-Lys, P11-Lys, and P11-N showed rare bactericidal activity against all Gram-negative and Gram-positive bacteria tested (Fig. 3c-d), which indicated that the antimicrobial peptides, namely P10-CP and P11-CP, were essential for the bactericidal activity of both enzymes. Curiously, for unknown reasons, P11-C showed higher bactericidal activity than the holoenzyme PHAb11 against all bacteria tested (Fig. 3d). Considering the different targets of antimicrobial peptides and peptidoglycan hydrolases, we further questioned the killing mechanism of PHAb10 and PHAb11 against susceptible bacteria. To this end, we evaluated the effects of different truncations of PHAb10 and PHAb11 on different bacteria by halo assay to test their peptidoglycan hydrolase activity. Results showed that lysozyme-containing truncations from both enzymes showed hydrolytic activity only towards peptidoglycan from Gram-negative bacteria but not from Gram-positive bacteria (Fig. 3e), suggesting that the killing of Gram-positive bacteria by PHAb10 and PHAb11 is probably mediated mainly by their internal antimicrobial peptides, i.e., P10-CP and P11-CP. Supporting to this assumption, rare peptidoglycan hydrolytic activity was observed in P10-CP and P11-CP (Fig. 3e). Altogether, these results suggested that the killing of Gram-negative bacteria by PHAb10 and PHAb11 might be a combination of lysozyme-mediated peptidoglycan hydrolytic activity and antimicrobial peptide-mediated outer membrane-disruption activity. However, their bactericidal activity against Gram-positive bacteria was primed solely by their internal antimicrobial peptides (Fig. 3f).

**Fig. 3.**
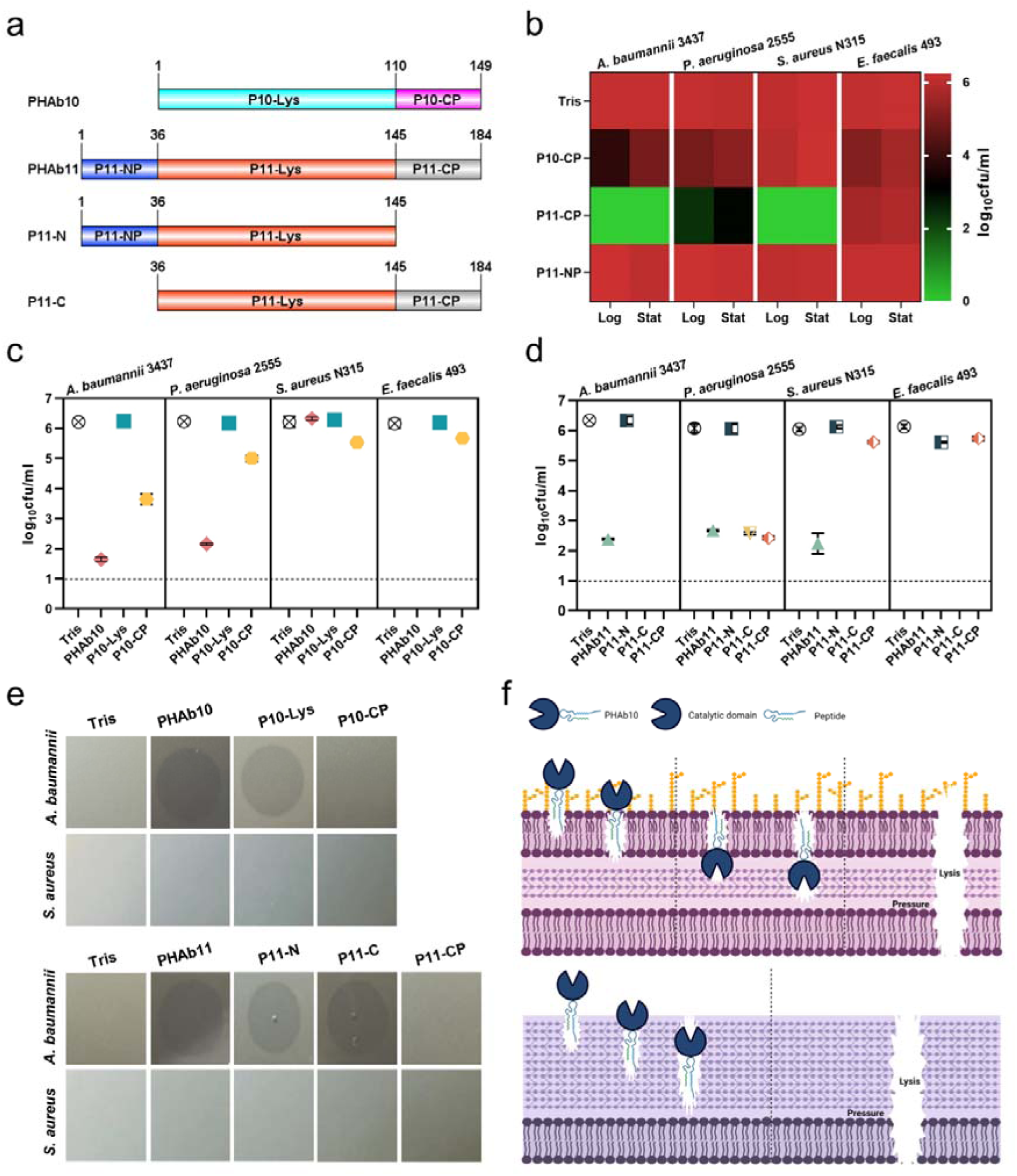
Mechanism of action of PHAb10 and PHAb11 on Gram-negative and Gram-positive bacteria. (a) Schematic representation of PHAb10 and PHAb11 domains. (b) Bactericidal activity of peptides derived from PHAb10 and PHAb11. Bacterial cells are treated with 50 μg/ml P10-CP, P11-NP, or P11-CP in 20 mM Tris-HCl at 37°C for 1 hour. (c-d) Bactericidal activity of different truncations of PHAb10 and PHAb11. Exponential *A. baumannii* 3437 cells are incubated with 50 μg/ml of each truncated fragment in 20 mM Tris-HCl for 1 hour at 37°C. Dash lines represent the limit of detection and data below the limit of detection is not shown. (e) Peptidoglycan hydrolytic activity of different truncations of PHAb10 and PHAb11. 0.1 μg of each truncated fragment is dropped onto autoclaved lawns of *A. baumannii* 3437 and *S. aureus* N315 and incubated at 37°C for 4 h. The groups treated with an equal volume of 20 mM Tris-HCl served as controls. (f) Schematic diagram of the bactericidal mechanisms of PHAb10 against Gram-negative bacteria (top) and Gram-positive bacteria (bottom).

### PHAb8 is a thermosensitive monomer, whilst PHAb10 and PHAb11 formed a thermostable dimer

To understand the catalytic mechanism and thermal stability of these peptidoglycan hydrolases, we attempted to decipher their structures and eventually obtained the crystal structures of PHAb8, PHAb10, and PHAb11 by X-ray crystallography (Table S2). Results showed that PHAb8 was a monomer (Fig. 4a and Fig. S4a), whilst PHAb10 (Fig. 4b) and PHAb11 (Fig. 4c) formed an asymmetric dimer. Specifically, the two asymmetric units of PHAb10 formed an antiparallel dimer with a rotation angle of 177.2° along the rotation axis and a distance of 10.5 Å between them. Further, PHAb10 dimer adopted a circularly permuted architecture, in which the N-terminus of one chain and the C-terminus of another chain formed a cavity, which might be the binding site of the peptidoglycan substrate (Fig. 4b). The two subunits of PHAb11 formed a tail-to-tail dimer with a rotation angle of −179.4° along the rotation axis and a distance of 37.1 Å between them. Each PHAb11 monomer contained an independent substrate-binding cavity around its catalytic triad (Fig. 4c). Notably, the N-terminal 36 amino acids, P11-NP, were missing in our PHAb11 dimer. We speculated that it may be destroyed during crystal formation. Interestingly, PHAb8 showed high topological similarity to monomeric PHAb10 and PHAb11, with the typical catalytic triad (Glu-Asp-Thr) located at the coil region (Fig. 4d and Fig. S4a-c). Dali-based screening further showed that the catalytic triad of PHAb8, PHAb10, and PHAb11 were highly conserved in homologous proteins, including AcLys (PDB ID 6ET6), LysF1 (PDB ID 7M5I), SpmX-Mur-*Ae* (PDB ID 6H9D), P22 (PDB ID 2ANX), and R21 (PDB ID 3HDE) (Fig. S4d and Table S3).

**Fig. 4.**
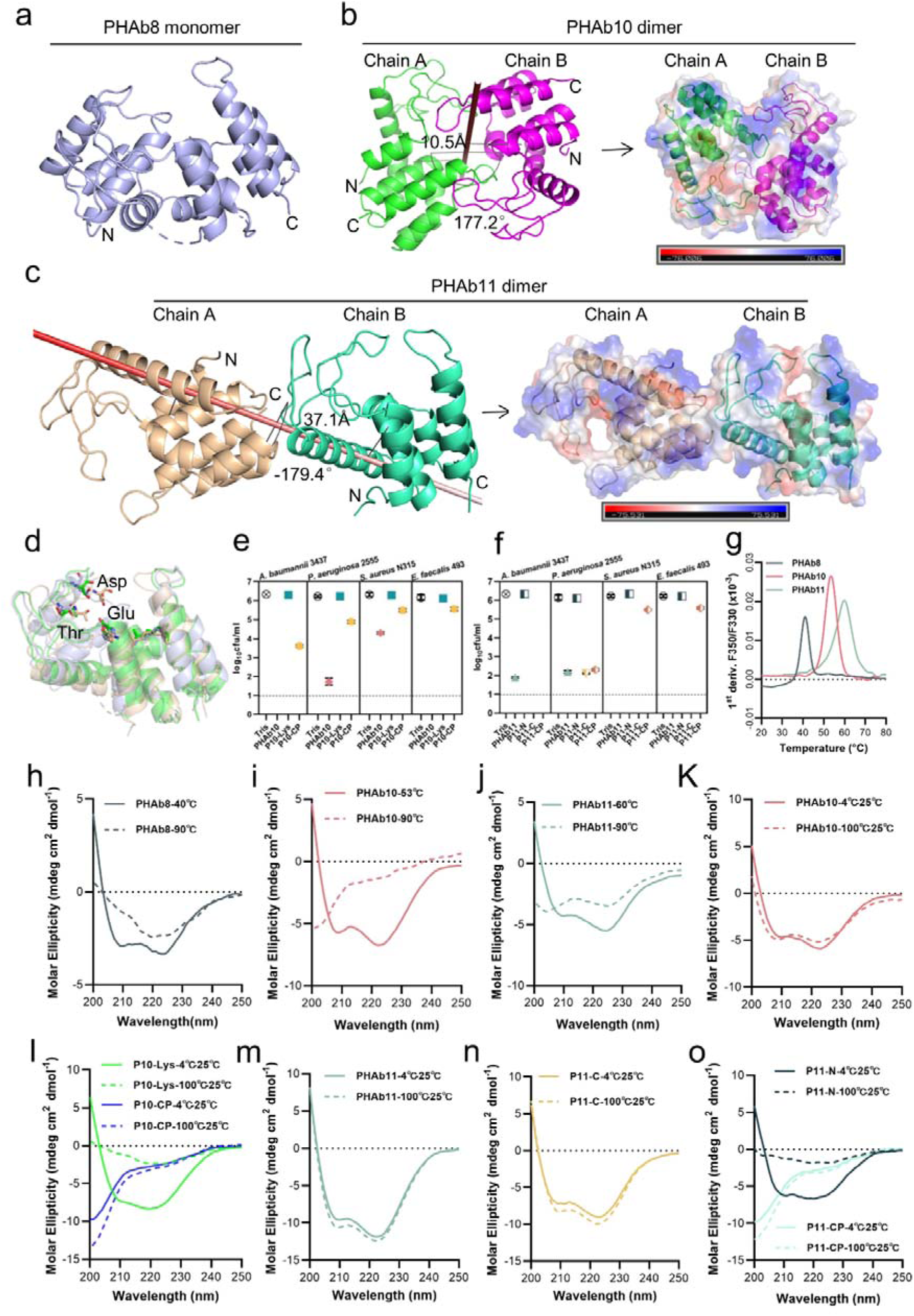
PHAb10 and PHAb11 are thermostable dimers. (a) Overall structure of PHAb8 monomer. (b) Dimeric structure of PHAb10. Chain A is colored in green, Chain B is colored in magenta, and the gray stick denotes the rotation axis. The electrostatic surface of PHAb10 with two potential substrate binding cavities are shown (negative = red; positive = blue). (c) Dimeric structure of PHAb11. Chain A is colored in wheat, Chain B is colored in green-cyan, and the red stick denotes the rotation axis. The electrostatic surface of PHAb11 with two potential substrate binding cavities is also shown (negative = red; positive = blue). (d) Superimposition of monomeric PHAb8, PHAb10, and PHAb11. PHAb8 is shown in light blue, PHAb10 in green, and PHAb11 in wheat. Residues involved in the catalytic triad are shown in sticks. (e-f) Residual bactericidal activity of PHAb10, PHAb11, and their truncation variants. Each truncation variant is treated at 100°C for 1 hour, stored at 25°C for 1 hour, and then tested for bactericidal activity by a log-killing assay. Dash lines represent the limit of detection and data below the limit of detection is not shown. (g) Thermal unfolding curves of PHAb8, PHAb10, and PHAb11 from 20 to 100°C as determined by nanoDSF. Values on the y-axis represent the first derivative of the fluorescence ratio at 350 nm and 330 nm. Peaks represent the transition temperature of each protein. (h-j) Circular dichroism spectra of PHAb8 (h), PHAb10 (i), and PHAb11 (j) at 90°C and temperatures close to their transition temperatures. (k-o) Circular dichroism spectra of PHAb10, PHAb11, and their truncation variants before and after heat treatment. Each domain fragment is treated at 100°C for 1 hour, stored at 25°C for additional hour prior to CD detection.

Superimposition of PHAb8, PHAb10, PHAb11 with other well-known lysozymes, including hen egg-white lysozyme (HEWL, PDB ID 4HPI), goose egg-white lysozyme (GEWL, PDB ID 153L), T4 lysozyme (T4L, PDB ID 1LYD), and lambda lytic transglycosylase (PDB ID 1LYD) revealed that the overall structures of PHAb8, PHAb10, and PHAb11 had relatively low similarity to HEWL, GEWL, and lambda lytic transglycosylase (Fig. S4e-g). Despite their lower amino acid sequence similarity (Fig. S4h), PHAb8, PHAb10, and PHAb11 superimposed well with T4 lysozyme, especially in their Glu-Asp-Thr catalytic triad regions (Fig. S4i and Table S3), suggesting that PHAb8, PHAb10, and PHAb11 may be phage-originated. Supporting to this, all three enzymes were predicted to be originated from prophages by PhageBoost (Fig. S5). All these observations together indicated that PHAb8, PHAb10, and PHAb11 were highly evolutionarily related to T4 lysozyme and might share similar catalytic mechanisms of action.

To reveal which domain contributes to the thermal stability of PHAb10 and PHAb11, we examined the residual bactericidal activity of their individual domains by treating them at 100°C for 1 hour. Results showed that for all strains tested, the holoenzyme and its antimicrobial peptide domain P10-CP retained high bactericidal activity, but its lysozyme catalytic domain P10-Lys did not (Fig. 4e). Except for the N-terminal peptide P11-NP, other truncated PHAb11 still maintained substantial bactericidal activity after heating at 100°C for 1 hour (Fig. 4f). These observations raised the question of whether PHAb10 and PHAb11 dimers are hyper-thermostable. To test this, we determined the temperature-dependent denaturation of these enzymes by nano-differential scanning fluorimetry (nanoDSF). Our results showed that PHAb8, PHAb10, and PHAb11 were denatured with transition temperatures of 41.4, 53.7, and 59.9°C, respectively (Fig. 4g). Interestingly, signs of aggregation were only observed in PHAb8 but not in PHAb10 and PHAb11 (Fig. S4j), indicating that PHAb10 and PHAb11 underwent structural refolding without precipitation during heat treatment. Consistent with these observations, the corresponding circular dichroism (CD) curves of PHAb8, PHAb10, and PHAb11 at 90°C were significantly different from their corresponding curves at temperature close to their transition temperature (Fig. 4h-j), suggesting that, similar to PHAb8, PHAb10 and PHAb11 lost their native conformation at high temperature such as 90°C. To further understand the thermodynamics of PHAb10 and PHAb11, we examined the structure of PHAb10 before and after heat treatment at 100°C for 1 hour by CD. Intriguingly, only minor differences were observed before and after heat treatment (Fig. 4k), indicating that PHAb10 almost refolded to its native conformation during cooling after heat treatment. The C-terminal antimicrobial peptide P10-CP of PHAb10 could withstand heat treatment, but P10-Lys was deformed after being treated at 100°C for 1 hour (Fig. 4l). Similar to the folding-refolding phenomenon of PHAb10, PHAb11 and its N-terminal peptide-deleted variant P11-C could restore to their native conformation after 1 hour treatment at 100°C (Fig. 4m-n), whilst its C-terminal peptide-deleted variant P11-N could not refold to its native conformation (Fig. 4o), indicating that C-terminal peptide was essential for its hyper-thermostability. Altogether, these observations suggested that dimerized PHAb10 and PHAb11 possessed unique folding-refolding thermodynamic mechanism.

### Folding-refolding thermodynamic of PHAb10 dimer is governed by intermolecular interactions

Since PHAb10 dimer and PHAb11 dimer exhibit a similar folding-refolding thermodynamic profiles, we wondered whether it was achieved through dimer-monomer switching. Therefore, we used PHAb10 as an example to test this hypothesis. We carefully examined the topology of the PHAb10 dimer and found that it was supported by 7 pairs of intermolecular H-bonds, with 3 pairs supporting the head, each 2 pairs in midbody and tail of the PHAb10 dimer (Table S4). Specifically, in the top view of chain A, Arg20 and Gly27 were involved in intermolecular H-bond interactions, supporting the head region, Gly128 and Gly129-mediated interaction contributed to the stability of midbody, whilst Lys133 and Arg137 were involved in intermolecular H-bond forces anchoring the tails of the PHAb10 dimer (Fig. 5a and Table S4). Next, we constructed several PHAb10 variants involving depletion of intermolecular force at the head, midbody, tail, or entire entity of the PHAb10 dimer (Table S5). Since PHAb10 was a verified dimer after chemical cross-linking in SDS-PAGE gel (Fig. 5b), we further examined the polymerization of these PHAb10 variants by Native-PAGE. As shown in Fig. 5c, wildtype PHAb10 showed two bands corresponding to its monomeric and dimeric forms. Whilst the variants R20IG27D, which has mutations in the head-supporting site, and G27DR137I, which has mutations in both the head and tail supporting sites, lost dimerization ability. As a result, both variants lost most of their bactericidal activity after treatment at 100°C for 1 hour (Fig. 5d), which could be further explained by the failure to recover their native forms after heat treatment (Fig. 5e-f). In support of these observations, the variant PHAb10-mut containing mutations in all 7 pairs of dimer-supporting sites showed aborted bactericidal activity (Fig. 5d) and failed to recover its native conformation (Fig. 5e-f) after treatment at 100°C for 1 hour. In contrast, the variants G128D/G129D with mutations in the midbody-supporting sites still showed a mixture of monomers and dimers (Fig. 5c), indicating that depletion of the midbody-supporting force hardly affected the dimerization of PHAb10. Consistent with this observation, the variant G128D/G129D still remained intact bactericidal activity (Fig. 5d) and could refold to its native form after heat treatment (Fig. 5e-f). Moreover, the variant K133A/R137I with mutations in the tail-supporting site also showed aborted bactericidal activity (Fig. 5d) and an irreversible conformation change after heat treatment (Fig. 5e-f), although it maintained a weak dimerization ability (Fig. 5c). This suggested that intermolecular forces at the tail of the PHAb10 dimer had only a small impact on its thermodynamics properties. Taken all these observations together, we proposed a model by which the PHAb10 dimer executed its folding-refolding thermodynamics (Fig. 5g). At temperatures above its transition point (i.e., >53.7°C), dimerized PHAb10 dissociated into monomers without precipitation. During the cooling process, monomeric PHAb10 refolded to a stable dimer *via* the intermolecular interactions between the two monomers, i.e., three pairs of intermolecular bonds at the head, two in the midbody, and two at the tail of the PHAb10 dimer. The folding and refolding dynamics of PHAb10 after heat treatment were somewhat similar to the opening and closing of zippers, where the intermolecular bonds act as inducers.

**Fig. 5.**
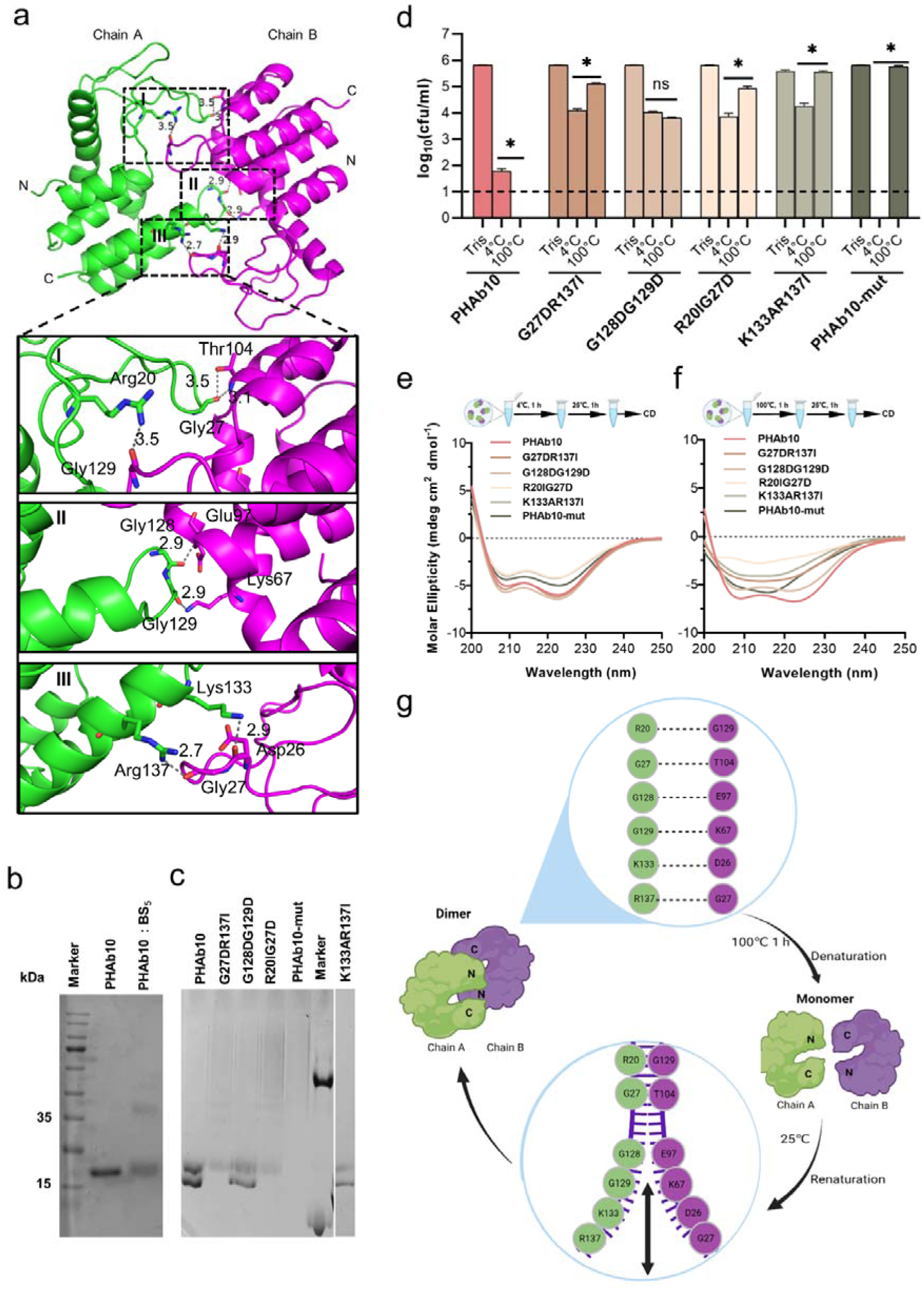
Folding-refolding thermodynamics of PHAb10. (a) Intermolecular interactions of PHAb10 dimer. The residues involved are shown as sticks. (b) SDS-PAGE analysis of PHAb10 with or without chemical cross-linker. (c) Native-PAGE analysis of PHAb10 and its variants. Marker: the upper band represents BSA (66.4 kDa), and the lower band represents lysozyme (14 kDa). (d) Bactericidal activity of PHAb10 and its variants before and after heat treatment. Each protein is incubated for 1 hour at 4 or 100°C, stored at 25°C for an additional 1 hour, and then examined for bactericidal activity in 20 mM Tris-HCl (pH 7.4) at a concentration of 50 μg/ml against exponential *A. baumannii* 3437 cells for 1 hour. Dash lines represent the limit of detection and data below the limit of detection is not shown. Data are analyzed by two-tailed Student’s t-tests. ns: statistically not significant; *: p<0.05. (e-f) Circular dichroism spectra of PHAb10 and its mutants before and after treatment at 100°C for 1 hour. (g) Zipper model showing the thermodynamics of PHAb10 dimer.

### PHAb10 efficiently removes *A. baumannii* from infection sites *in vivo*

Since PHAb10 exhibited rare cytotoxicity at the cellular level (Fig. S6), we further tested its bactericidal activity in two different mouse infection models. In a burn wound model of *A. baumannii* infection (Fig. 6a), mice were first treated with 10 µg/mouse PHAb10 (10 µl; n = 5) for 4 hours, and then infected with 10 µl 5×10^8^ cfu/ml of *A. baumannii* 3437 for 24 hours. We found significantly less viable bacteria remaining at the site of infection compared to Tris-treated controls (Fig. 6b). Treatment with 4 µg/mouse minocycline (10 µl; n = 5) also showed good bacterial removal compared to the control group, however, no statistical difference was observed between groups treated with 10 µg/mouse PHAb10 and 4 µg/mouse minocycline (Fig. 6b), suggesting that when administered topically, PHAb10 was as efficient as minocycline in clearing susceptible bacteria from the site of infection. In a mouse abscess model, mice were infected hypodermically (i.h.) with 25 μl of *A. baumannii* 3437 at a concentration of 5×10^8^ cfu/ml for 24 hour, and then treated with 10 µg/mouse PHAb10 (10 µl; n = 6) or 4 µg/mouse minocycline (10 µl; n = 7) subcutaneously for 5 days. Six days after infection, viable bacteria were recovered from the infection site and counted (Fig. 6c). Our results showed that there was rare difference between the minocycline-treated group and the Tris-treated control group (Fig. 6d). In contrast, a significant reduction in residual *A. baumannii* was observed in the PHAb10-treated group (Fig. 6d), indicating that PHAb10 had better bactericidal efficacy than minocycline when injected subcutaneously at the site of infection in a skin infection model. Notably, there is little difference in the change in animal body weight between groups treated with PHAb10, minocycline, or Tris buffer (Fig. S7). Taking all these observations together, PHAb10 appears to exhibit good bacterial clearance efficacy *in vivo* and may represent a potential antimicrobial agent for the treatment of drug-resistant bacteria.

**Fig. 6.**
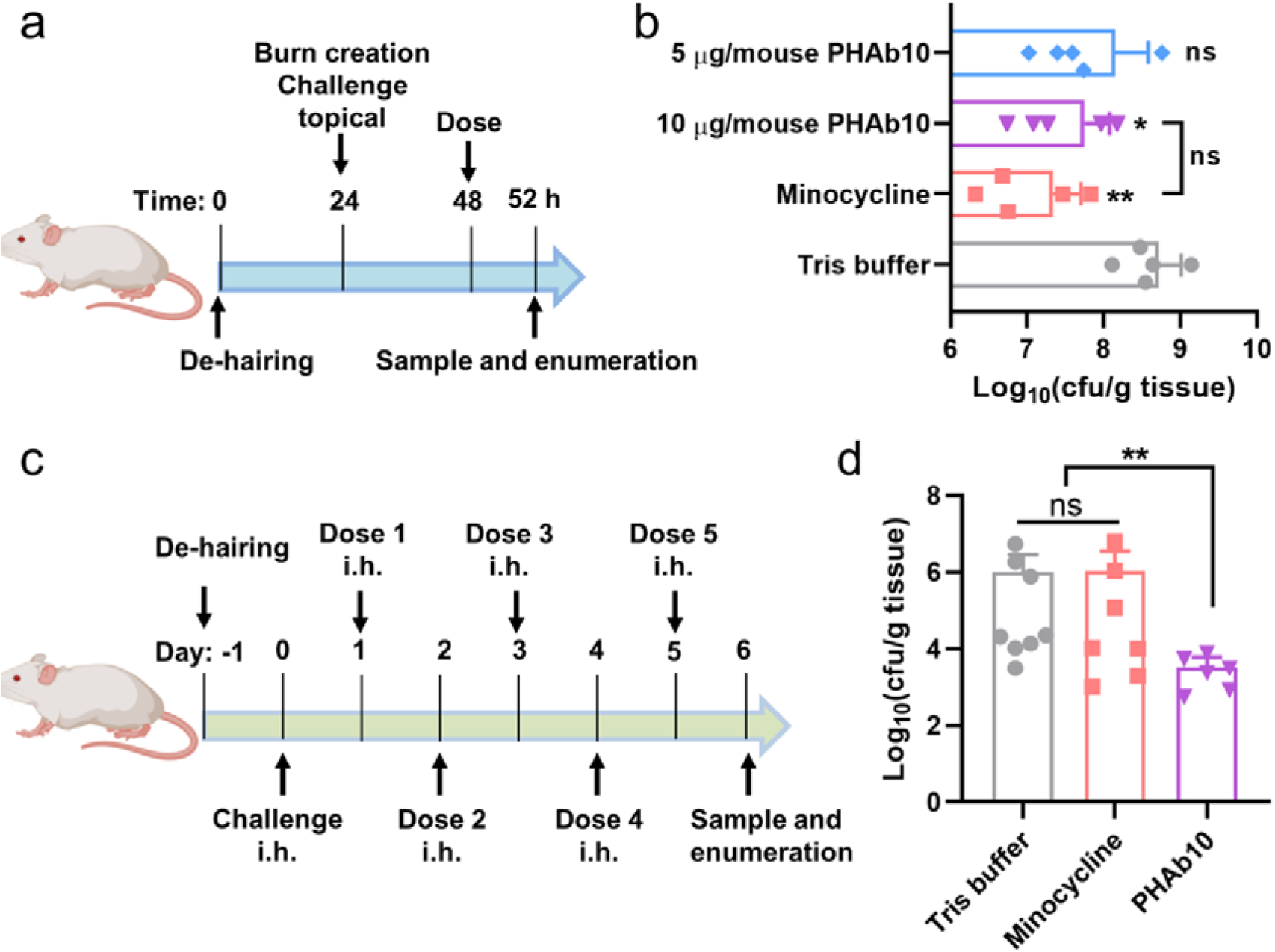
PHAb10 shows high efficacy in mouse infection models. (a) Experimental schema for burn wound infection model. Burn wounds produced by exposing naked skin to 65°C water for 12 s was infected with 10 µl 5×10^8^ cfu/ml of *A. baumannii* 3437. At 24-hour post-colonization, mice are treated with 5 or 10 µg/mouse PHAb10 (10 µl; n = 5), 4 µg/mouse minocycline (10 µl; n = 5), or an equal volume of Tris buffer (10 µl; n = 5). Four hours post-treatment, viable bacteria on the burn wound skin is collected and counted. (b) Viable *A. baumannii* collected from burn wound skin after each treatment. The number of viable cells in each group is normalized and compared with the number of viable cells treated with Tris buffer by one-way analysis of variance (ANOVA). ns: statistically not significant; *: p<0.05; **: p<0.01. (c) Experimental schema for abscess model. Mice are infected hypodermically with 25 μl 5×10^8^ cfu/ml of *A. baumannii* 3437 on the right side of the back dorsum. Twenty-four hours after infection, mice are injected hypodermically with 10 µg/mouse PHAb10 (10 µl; n = 6), 4 µg/mouse minocycline (10 µl; n = 7), or an equal volume of Tris buffer (10 µl; n = 8). Each group is injected subcutaneously once a day for 5 consecutive days. Viable bacteria were counted at the infection site 24 hours after the last dose. (d) Viable *A. baumannii* collected from the skin after each treatment. The number of viable cells from each group was normalized and compared with those of the Tris buffer-treated group by ANOVA. ns: statistically not significant; **: p<0.01.

## Discussion

The uncontrollable spread and emergence of antimicrobial resistance requires the discovery of new antimicrobial agents with different targets or mechanisms of action than traditional antibiotics. Bacteriophage-derived peptidoglycan hydrolases (i.e., lysins) have been investigated as promising alternatives to combat resistant bacteria due to their unique mechanism of action, potent bactericidal activity, low risk of drug resistance development. However, for Gram-negative bacteria, the effects of phage-derived lysins were often hampered by their outer membranes, which requires more strategies to overcome this additional barrier. Although some engineered lysins (especially Artilysins, Innolysins, and Lysocins) have shown promising performance, clinically promising lysins capable of destroying Gram-negative bacteria remain inadequately addressed. Considering the fact that lysins from Gram-negative bacteria usually contain antimicrobial peptide-like regions (Vazquez, Garcia et al. 2021), in the present work, we report a new strategy to mine active bactericidal peptidoglycan hydrolases from bacterial proteomes through lysin-derived antimicrobial peptide-priming screening. As a proof-of-concept, we successfully discovered the two peptidoglycan hydrolases PHAb10 and PHAb11 from the proteome of *A. baumannii* using the PlyF307 lysin-derived peptide as a template, highlighting the utility of this strategy in mining antimicrobial agents from bacterial database.

Notably, PHAb10 and PHAb11 exhibited high bactericidal activity against a variety of Gram-negative and Gram-positive microbes. Unlike other Gram-negative lysins that act in an outer membrane permeabilizer-dependent manner (Briers, Walmagh et al. 2011, Walmagh, Boczkowska et al. 2013, Oliveira, Thiagarajan et al. 2014, Plotka, Kapusta et al. 2019), PHAb10 and PHAb11 exhibited intrinsic activity against Gram-negative and Gram-positive pathogens *via* different mechanisms of action. Due to their C-terminal antimicrobial peptides P10-CP and P11-CP, PHAb10 and PHAb11 can penetrate the outer membrane of Gram-negative bacteria in a self-primed manner, allowing them to enter and cleave peptidoglycan bonds, ultimately leading to osmotic lysis of Gram-negative pathogens, while Gram-positive bacteria are killed mainly through the action of the intrinsic antimicrobial peptides.

Structure-based homology analysis revealed that PHAb10 and PHAb11 probably belonged to the T4 lysozyme family and possessed the conserved Glu-Asp-Thr catalytic triad shared by traditional phage lysins and bacterial autolysins, namely Glu17, Asp26, and Thr32 of PHAb10, and Glu52, Asp61, and Thr67 of PHAb11. It is known that T4 lysozyme (T4L) from T4 phage completely loses its enzymatical activity after treatment at 75°C for 5 minutes (Tsugita, Inouye et al. 1968), but somehow, heat-denatured T4L retains its bactericidal activity because of the amphipathic α-helix region at the C-terminus of the enzyme which contains membrane disturbing activity (During, Porsch et al. 1999). Unlike T4L, whose monomer crystals have been exhaustively studied, PHAb10 or PHAb11 is a homodimer that retain almost 100% of their bactericidal activity after heat treatment through a unique dimer-monomer transition. Interestingly, several T4L mutants have been reported to be homodimers crystallographically (Banatao, Cascio et al. 2006), although naturally occurring dimeric T4L-like enzymes are still rarely reported. Notably, heat treatment converts hen egg white lysozyme (HEWL) to a partially unfolded, enzymatically inactive and more hydrophobic dimeric form, which exhibits an enhanced bactericidal activity against Gram-negative bacteria (Ibrahim, Higashiguchi et al. 1996). The excellent thermostability capacity of PHAb10 and PHAb11 distinguishes them from most other enzymes reported to date. For example, gp36C lysin from bacteriophage [KMV, a known highly thermostable lysozyme, retains only [50% residual activity after treatment at 100°C for 1 hour (Lavigne, Briers et al. 2004, Briers, Lavigne et al. 2006). Heat-resistant lysin have potential applications in animal feed premixes (pelletizing temperature 75-98°C) and medical devices requiring autoclaving.

To the best of our knowledge, few proteins have been demonstrated to have reversible folding-refolding thermodynamics at high temperature, but their mechanisms are unclear, such as HPL118, HPL511, and HPLP35 (Schmelcher, Waldherr et al. 2012). Here we found that PHAb10 had unique temperature-dependent folding-refolding thermodynamic, and structural and biochemical analyses further suggested that PHAb10 undergoes a unique dimer-monomer transition mediated by 7 pairs of intermolecular interactions to survive thermal treatment.

Bacteriophages are known to be the most abundant biological entities on earth (Knowles, Silveira et al. 2016). Over billions of years of coevolution, more than 80% of bacterial genomes have become populated with at least one prophage (Keen and Dantas 2018, Miller-Ensminger, Garretto et al. 2018), also known as molecular imprints, and which can serve as an untapped arsenal for the discovery of phage-related antimicrobial agents. Notably, prophage identification lags greatly behind genome or metagenome sequencing. Therefore, there is reason to believe that advances in bacterial culturomics (Lagier, Dubourg et al. 2018), ‘rebooting’ phage genomes in more tractable hosts (Kilcher, Studer et al. 2018), and computational biology (Arndt, Marcu et al. 2019, Song, Sun et al. 2019), may help discover otherwise inaccessible prophages from bacterial genome big data. In turn, this will increase the added value of bacterial genomes as a new source of unearthed therapeutically promising antimicrobial drugs. More importantly, our findings implied that big data, such as bacterial genomes and proteomes, human proteomes and other metagenomic datasets, could in principle become an encrypted power in the fight against antimicrobial resistance.

## Materials and Methods

### Bacterial strains

The bacterial strains used in this work are described in Table S6. *Acinetobacter baumannii*, *Escherichia coli*, *Pseudomonas aeruginosa*, *Klebsiella pneumoniae*, *Enterococcus faecalis,* and *Staphylococcus aureus* were all grown in Lysogeny Broth (LB) at 37°C. *Streptococcus pneumoniae* was statically cultured in Todd Hewitt Broth with 0.5% Yeast extract (THY) at 37°C with 5% CO_2_. All other *Streptococci* were grown in Brain Heart Infusion (BHI) broth at 37°C.

### Screening of putative peptidoglycan hydrolases from *A. baumannii*

Putative peptidoglycan hydrolases from the *A. baumannii* (PHAbs) proteome database were obtained by BLASTP in NCBI using the lysin PlyF307-derived antimicrobial peptide P307 as input sequence. Sequences marked as partial, shorter than 110 amino acids, longer than 300 amino acids, or clearly not belonging to lysins were manually excluded from further analysis.

Multiple sequence alignments were performed by the ClustalW algorithm in MEGAX software and visualized by Jalview. Phylogenetic trees were constructed by MEGAX software using the Neighbor-Joining method and visualized by the iTOL online service (http://itol.embl.de/). Three-dimensional (3D) protein models showing different amino acid residues were predicted online by SWISS-MODEL (https://swissmodel.expasy.org/).

### *In silico* analysis of PHAbs

The basic physical and chemical properties of PHAbs were analyzed by ProtParam (https://web.expasy.org/protparam/). Solubility was predicted by Protein-Sol (https://protein-sol.manchester.ac.uk/), and signal peptides were predicted by SignalP (https://services.healthtech.dtu.dk/service.php?SignalP-5.0). The transmembrane area was predicted by TMHMM (https://services.healthtech.dtu.dk/service.php?TMHMM-2.0), while domain predictions were performed through CD searches in the NCBI database (https://www.ncbi.nlm.nih.gov/Structure/cdd/wrpsb.cgi) and InterPro in the EBI database (http://www.ebi. ac.uk/interpro/search/sequence/). The Percent Identity Matrix was calculated by Clustal Omega (https://www.ebi.ac.uk/Tools/msa/clustalo/) and visualized with TBtools. The prophage origination analysis was predicted by PhageBoost (https://phageboost.ku.dk/) (Siren, Millard et al. 2021). The net charge of the N-terminal and C-terminal 30 amino acids of each PHAb was calculated by BaAMPs (http://www.baamps.it/tools/calculator) under neutral condition. The Local Net Charge of each PHAb was obtained by assigning 1 to the basic amino acids Arg (R) and Lys (K) and −1 to the acidic amino acids Glu (E) and Asp (D) and summing all assignments in a 31 amino acid window composing of 15 amino acids before and after the determined position. The 3D models of each PHAb were predicted by AlphaFold2 and visualized by PyMOL.

### Gene synthesis and cloning

The coding sequences of the original five putative peptidoglycan hydrolases, namely, PHAb7, PHAb8, PHAb9, PHAb10, and PHAb11, as well as the PHAb10 variant, PHAb10-mut with 6 site mutations were chemically synthesized with codon optimization and cloned into a pET28b(+) vector by *Nco*I and *Xho*I restriction sites. Additional variants of PHAb10 and truncations of PHAb10 and PHAb11 were constructed by primer-derived overlap PCR (Table S7). All constructs were electro-transformed into *E. coli* BL21(DE3) competent cells for expression. Peptides derived from PHAb10 and PHAb11, namely P10-CP (amino acids 111-149), P11-NP (amino acids 1-36), and P11-CP (amino acids 146-184), were chemically synthesized and dissolved in ultrapure water before use.

### Protein purification

BL21(DE3) cells were grown in LB to an optical density at 600 nm (OD_600_) of 0.4-0.6 and induced with 0.2 mM isopropyl β-D-thiogalactopyranoside at 37°C for 2 h. Cells were then harvested and lysed by sonication. Supernatant was collected and passed through a Ni-NTA column pre-equilibrated with 20 mM imidazole. Fractions were collected by washing and eluting with 60 mM and 250 mM imidazole, respectively. The collected active protein fractions were pooled, dialyzed against 20 mM Tris-HCl (pH 7.4), and then filter-sterilized. Protein for crystallization was further purified by size-exclusion chromatography on a Superdex 75 (GE Healthcare) with gel filtration buffer (50 mM Tris-HCl, pH 8.0, 150 mM NaCl, 1 mM DTT). Protein purity was examined by 12% SDS-PAGE, and protein concentration determined using the BCA Protein Concentration Assay Kit. Dimerization of PHAb10 and its variants was detected by Native-PAGE.

### Halo assay

Bacterial strains were grown overnight in 100 ml LB, harvested, and resuspended in 50 ml phosphate buffered saline (PBS) containing 0.7% agarose. Cells were then autoclaved and poured into glass Petri dish culture plates. Ten microliter of peptidoglycan hydrolases at different concentrations (0, 2.5, 5, and 10 μg/ml) were dropped onto each bacterial lawn and incubated at 37°C till a clear zone was formed, and its area was determined using the method previously described (Zhang, Cai et al. 2017, Vander Elst, Linden et al. 2020). Wells treated with an equal volume of 20 mM Tris-HCl were used as controls.

### Antibacterial activity assay

To determine the antibacterial activity of peptidoglycan hydrolases, bacteria were cultured overnight (stationary phase, OD_600_ = 1.2-1.6), then transferred to fresh medium at 100-fold dilution and continued to grow for 3 hours to OD_600_ = 0.5-0.6 in the exponential growth phase. Cultures of different bacteria were centrifuged at 10,000 × *g* for 1 minute, washed once and resuspended in 20 mM Tris-HCl, pH 7.4. Stationary phase bacterial cells were diluted to a final OD_600_ of 0.6 before use. To test the bactericidal activity of each peptidoglycan hydrolase, 100 μl of a bacterial suspension in stationary phase or exponential phase was mixed with an equal volume of peptidoglycan hydrolase at different concentrations at 37°C for 1 hour with shaking at 200 rpm. The number of viable bacteria after each treatment was counted by plating serial dilutions on LB agar. Wells treated with an equal volume of dialysis buffer instead of peptidoglycan hydrolases were used as controls. Specifically, to test the dose-dependent antibacterial activity of PHAb10 and PHAb11, exponential cultures of various Gram-negative and Gram-positive bacteria were treated with 0, 0.625, 1.25. 2.5, 5, 6.25, 10, 12.5, 25, 50, or 100 μg/ml of each enzyme in 20 mM Tris-HCl (pH 7.4) at 37°C for 1 hour.

To test the time-killing curves of PHAb10 and PHAb11 against exponential *A. baumannii* 3437, bacterial suspensions were incubated with 50 μg/ml of each peptidoglycan hydrolase in 20 mM Tris-HCl (pH 7.4) for 0, 0.5, 1, 3, 5, 10, 15, 30, 45, 60, and 120 min at 37°C.

To measure the effects of different environmental factors on the antibacterial activity of PHAb10 and PHAb11, exponential *A. baumannii* 3437 cells were treated with 50 μg/ml of each peptidoglycan hydrolase at different pH values (5, 6, 7, 8, 9, and 10), various temperatures (4, 25, 37, 45, 55, 65, 75, 85, and 100°C), and various NaCl concentrations (0, 25, 50, 100, 200, 500, and 1000 mM), and urea (0, 50, 100, 200, 500, and 1000 mM) at 37°C for 1 hour. All assays were performed at least three times in biological replicates.

### Thermostability assay

The thermostability of PHAb and their variants at 100 μg/ml was determined by a Prometheus NT.48 nano-differential scanning fluorimetry (nanoDSF, NanoTemper Technologies, CA, USA) in the temperature range of 20-100°C (increasing step of 1°C/min) in 20 mM Tris-HCl (pH 7.4). The first derivative of the fluorescence ratio at 350 and 330 nm (1st der, F350/F330) was calculated automatically by the PR-ThermControl software supplied with the instrumentation. The transition temperature (Tm) corresponds to the 1st order peak of F350/F330.

### Circular dichroism

The circular dichroism spectra of PHAb and its variants at 200 µg/ml in 20 mM Tris-HCl (pH 7.4) were collected by an Applied Photophysics Chirascan Plus circular dichroism spectrometer (Leatherhead, UK) from 200-260 nm (0.1 cm path length) at room temperature.

The spectra of air and Tris-HCl buffer (pH 7.4) were recorded as background and baseline, respectively. Secondary structures were calculated by CDNN V2.1 software supplied by the instrument manufacturer.

### Crystallization, data collection and processing

In this study, crystals were obtained for each protein at a concentration of 20-25 mg/ml using sitting drop method at a 1:1 ratio on a reservoir of 0.1 M citric acid pH 3.5, 25% w/v polyethylene glycol 3350 after 1 week of incubation at 16°C. They were then immersed in a cryoprotectant consisting of a reservoir solution supplemented with 10% glycerol before being flash frozen in liquid nitrogen. Diffraction data were collected at beamline 19U1 (BL19U1) at Shanghai Synchrotron Radiation Facility (SSRF), and the X-ray datasets were processed using HKL2000 and XDS software programs. The initial phase information was determined by molecular replacement in the program PHASER using *A. baumannii* AB 5075UW prophage (PDB code: 6ET6) as the initial search model. The structure was then improved by multiple rounds of manual construction and refinement by COOT and PHENIX. The final structures of PHAb8, PHAb10, and PHAb11 were visualized using PyMOL. Summary of data collection and refinement statistics are presented in Table S2.

### Cytotoxicity testing

The cytotoxicity of PHAb10 against HepG2 cells was determined by a Cell Counting Kit-8 (CCK-8) assay (Dojindo Molecular Technologies, Kumamoto, Japan) according to the manufacturer’s protocol. HepG2 cells were seeded in 96-well plates at a density of 5,000 cells per well in Dulbecco’s modified Eagle medium (DMEM; Sigma-Aldrich, Shanghai, China) supplemented with 10% fetal bovine serum, 1% penicillin, and 1% streptomycin for 24 h. The cells were then exposed to a series of concentrations of PHAb10 (0, 6.25, 12.5, 25, 50, and 100 μg/ml) for another 24 h. Afterwards, the contents of the plates were replaced with fresh medium containing 10% CCK-8 solution and incubated at 37°C for 1.5 h. The final optical density at OD450 was noted by use of a microplate reader (SynergyH1; BioTek, USA). The results were expressed as relative cell viability, expressed as a percentage of the growth of cells in control wells treated with PBS only.

### Mouse experiments

All mouse infection experiments were conducted in an ABSL-2 lab, and all experimental methods were carried out in accordance with the regulations and guidelines set forth by the Animal Experiments Committee of the Wuhan Institute of Virology, Chinese Academy of Sciences. All experimental protocols were approved by the Animal Experiments Committee of Wuhan Institute of Virology, Chinese Academy of Sciences (WIVA17202102). During the experiment, animals were housed in individually ventilated cages following a range of animal welfare and ethical criteria and were euthanized at the end of observation. In a burn infection model, 6-to 8-week-old female BALB/c mice were anesthetized, shaved, and depilated to create a 2-cm^2^ hairless area on the dorsum as previously described (Li, Jiang et al. 2021). Partial thickness burn was achieved by exposing bare skin to water at 65°C for 12 s. The burns were then inoculated with 10 µl 5×10^8^ cfu/ml of *A. baumannii* 3437. 24 hours after colonization, mice were randomly separated into four groups and treated with 5 or 10 µg/mouse PHAb10 (10 µl; n = 5), 4 µg/mouse minocycline (10 µl; n = 5), or an equal volume of Tris buffer (10 µl; n = 5). Four hours after treatment, mice were sacrificed by cervical dislocation to obtain skin. Specifically, the infected skin was excised, infiltrated in 1ml PBST (PBS containing 0.1% Triton X-100) for 2 min, and homogenized using a NewZongKe MD1000 Tissue Cell-Destroyer (NewZongKe, Wuhan, Hubei, China). The resulting solution was serially diluted and plated on LB agar containing 4 µg/ml gentamicin and 2 µg/ml meropenem.

In a mouse abscess model, 6-to 8-week-old female mice were depilated to create a 2-cm^2^ hairless area on the dorsum as described above and infected hypodermically (i.h.) with 25 μl of *A. baumannii* 3437 at a concentration of 5×10^8^ cfu/ml on the right side of the back dorsum. After 24 h of infection, mice were randomly divided into 3 groups and hypodermically treated with 10 µg/mouse PHAb10 (10 µl; n = 6), 4 µg/mouse minocycline (10 µl; n = 7), or an equal volume of Tris buffer (10 µl; n = 8). Subcutaneous dosage was performed once a day for 5 consecutive days for each group. The body weight of mice in each group was monitored daily. Twenty-four hours after the last dosage, mice were sacrificed by cervical dislocation. Infected skin was excised, infiltrated in 1ml PBST for 2 min, and subjected to tissue homogenization using a NewZongKe MD1000 Tissue Cell-Destroyer. Number of viable bacteria in each group was enumerated by plating serial dilutions on LB agar containing 4 µg/ml gentamicin and 2 µg/ml meropenem.

### Statistical analysis

Two-tailed Student’s t-tests were used to analysis all *in vitro* assays. In mouse models, viable bacterial cell numbers in each group were compared and analyzed by one-way analysis of variance (ANOVA). ns: statistically not significant; *: p<0.05; **: p<0.01.

## Data availability

The atomic coordinate and structure factors of PHAb8, PHAb10, and PHAb11 have been deposited in the PDB database with accession numbers 8HEM, 7YKU, and 8HDQ, respectively.

## Acknowledgments

This work was supported by the National Natural Science Foundation of China (No. 32070187, 32161133003, 31770192, and 81802001), and the Open Research Fund Program of National Bio-Safety Laboratory, Wuhan (No. 2021SPCAS001). We thank the staff of BL19U1 beamline at National Center for Protein Sciences Shanghai and Shanghai Synchrotron Radiation Facility (Shanghai, China) for assistance during data collection. We thank Dr. Xuefang An, and Dr. Yanfeng Yao from the Core Facility and Technical Support, Wuhan Institute of Virology for their assistance in animal experiments.

## Author contributions

**Li Zhang**: Investigation; Methodology; Writing—review and editing. **Fen Hu**: Investigation; Methodology; Funding acquisition; Writing—original draft; Writing—review and editing. **Zirong Zhao**: Investigation; Methodology; Writing—review and editing. **Xinfeng Li**: Investigation; Methodology; Writing—review and editing. **Mingyue Zhong**: Investigation; Methodology; Writing—review and editing. **Jiajun He**: Investigation; Methodology, Writing—review and editing. **Fangfang Yao**: Investigation, Methodology; Writing—review and editing. **Xiaomei Zhang**: Methodology; Writing—review and editing. **Yuxuan Mao**: Methodology; Writing—review and editing. **Hongping Wei**: Supervision; Methodology; Writing—review and editing. **Jin He**: Conceptualization; Supervision; Methodology; Writing—review and editing. **Hang Yang**: Conceptualization; Supervision; Investigation; Methodology; Methodology; Funding acquisition; Writing—original draft; Writing—review and editing.

