## Supplemental Table 1-7 and Supplemental Figure 1-7. for "Dimer-monomer transition defines a hyper-thermostable peptidoglycan hydrolase mined from bacterial proteome by lysin-derived antimicrobial peptide-primed screening"

**Table S1. Physicochemical properties of putative antimicrobial peptides in PHAb10 and PHAb11**

| Parameter | Amino acid sequence | Molecular weight (kDa) | pI | Negatively /positively charged residues | Net charg e | Instability index | Aliphatic index | GRAVY |
|---|---|---|---|---|---|---|---|---|
| P10-CP | NKGDYQ GAADQFL VWNKAG GKVMKGL VRRREAE RALFLKK | 4550 | 10.3 7 | 4/10 | 6 | 39.66 | 73.25 | -0.733 |
| P11-CP | NKGDYKG AADQFLV WNKAGG KVLKGLV RRREAER ALFLKK | 4532 | 10.4 5 | 4/11 | 7 | 35.41 | 83.00 | -0.695 |
| P11-NP | MTTKPFF DAARVIA GGKLTQA QVDELNK VVNKLAP G | 3829 | 9.52 | 3/5 | 2 | 19.74 | 89 | -0.053 |

Positively charged amino acids are labeled in red.

GRAVY: Grand average of hydropathicity.

**Table S2. Data collection and refinement statistics of PHAb8, PHAb10, and PHAb11 structures**

| Parameters | PHAb10 | PHAb11 | PHAb8 |
|---|---|---|---|
| **Data collection** | | | |
| Wavelength (Å) | 0.9793 | 0.9793 | 0.9793 |
| Space group | $P\ 1\ 2_1\ 1$ | $P\ 2\ 2\ 2_1$ | $C\ 2\ 2\ 2_1$ |
| Cell dimensions | | | |
| a, b, c (Å) | 31.86 73.77 78.62 | 50.01 54.9 161.31 | 79.03 156.98 47.26 |
| α, β, γ (°) | 90.00 93.02 90.00 | 90 90 90 | 90 90 90 |
| Resolution range (Å) | 31.82 - 1.52 (1.574 - 1.52) | 36.62 - 2.27 (2.351 - 2.27) | 32.06 - 2.4 (2.486 - 2.4) |
| Unique reflections | 55245 (5468) | 21246 (2049) | 11880 (1048) |
| Multiplicity | 6.8 (6.5) | 12.3 (12.4) | 12.9 (13.2) |
| Completeness (%) | 98.86 (98.17) | 99.65 (99.80) | 94.57 (88.22) |
| Mean I/sigma I | 18.36 (1.67) | 14.54 (2.02) | 10.50 (1.53) |
| Wilson B-factor ($Å^2$) | 24.27 | 31.87 | 37.86 |
| $R_{merge}$ | 0.04816 (1.097) | 0.2869 (1.486) | 0.213 (1.914) |
| $R_{meas}$ | 0.0522 (1.192) | 0.2999 (1.55) | 0.2219 (1.991) |
| $CC_{1/2}$ | 0.999 (0.788) | 0.992 (0.747) | 0.996 (0.579) |
| **Refinement** | | | |
| Resolution (Å) | 31.82 - 1.52 | 36.62 - 2.27 | 32.06 - 2.4 |
| $R_{work}$ | 0.1921 (0.3344) | 0.2483 (0.3092) | 0.2520 (0.3560) |
| $R_{free}$ | 0.2151 (0.3653) | 0.2660 (0.3670) | 0.2849 (0.4396) |
| Ligands | 0 | 18 | 17 |
| Rmsd from ideal | | | |
| Bond lengths (Å) | 0.004 | 0.005 | 0.010 |
| Bond angles (°) | 0.62 | 0.75 | 1.18 |
| Ramachandran plot | | | |
| Favored (%) | 98.97 | 98.25 | 95.81 |
| Allowed (%) | 1.03 | 1.75 | 4.19 |
| Outliers (%) | 0.00 | 0.00 | 0.00 |
| Average B-factor | 32.18 | 37.23 | 52.84 |

| PDB Code | 7YKU | 8HDQ | 8HEM |
| --- | --- | --- | --- |

Statistics for the highest-resolution shell are shown in parentheses.

**Table S3. T4L-like lysozymes with characterized structures**

| Protein | Catalytic Residues | | | PDB ID |
|---|---|---|---|---|
| PHAb8 | Glu74 | Asp83 | Thr89 | 8HEM |
| PHAb10 | Glu17 | Asp26 | Thr32 | 7YKU |
| PHAb11 | Glu52 | Asp61 | Thr67 | 8HDQ |
| AcLys | Glu64 | Asp73 | Thr79 | 6ET6 |
| LysF1 | Glu15 | Asp24 | Thr30 | 7M5I |
| SpmX-Mur-*Ae* | Glu18 | Leu27 | Met33 | 6H9D |
| P22 | Glu16 | Asp25 | Thr31 | 2ANX |
| R21 | Glu35 | Asp44 | Thr50 | 3HDE |
| HEWL | Glu35 | Asp52 | | 4HPI |
| GEWL | Glu73 | | | 153L |
| phage lambda lytic transglycosylase | Glu19 | | | 1D9U |
| T4L | Glu11 | Asp20 | Thr26 | 1LYD |

**Table S4. Intermolecular interactions of PHAb10 dimer**

| PHAB10 dimer | Serial number | Donor (atom 1) | Acceptor (atom 2) | Distance/Å | Force |
|---|---|---|---|---|---|
| Head-supporting force | 1 | R20.NH1 (Chain A) | G129.O (Chain B) | 3.5 | H-bond |
| | 2 | T104.N (Chain B) | G27.O (Chain A) | 3.1 | H-bond |
| | 3 | T104.OG1 (Chain B) | G27.O (Chain A) | 3.5 | H-bond |
| Midbody-supporting force | 4 | E97.N (Chain B) | G128.O (Chain A) | 2.9 | H-bond |
| | 5 | K67.NZ (Chain B) | G129.O (Chain A) | 2.9 | H-bond |
| Tail-supporting force | 6 | K133.NZ (Chain A) | D26.O (Chain B) | 2.9 | H-bond |
| | 7 | R137.NH1 (Chain A) | G27.O (Chain B) | 2.7 | H-bond |

**Table S5. PHAb10 variants**

| Native amino acids (Chain A) | Substituted amino acids | Mutant name |
|---|---|---|
| R20, G27 | R20I, G27D | R20I/G27D |
| G128, G129 | G128D, G129D | G128D/G129D |
| G27, R137 | G27D, R137I | G27D/R137I |
| K133, R137 | K133A, R137I | K133A/R137I |
| R20, G27, K67, G128, K133, R137 | R20E, G27T, K67W, G128N, K133A, R137N | PHAb10-mut |

**Table S6 Bacterial strains used in this work.**

| **Strain** | **Source** | **Strain** | **Source** |
|---|---|---|---|
| ***Acinetobacter baumanni*** | | ***Klebsiella pneumoniae*** | |
| ABA3437 | WIV[b] | KP21 | WIV[b] |
| CRAB2 | WIV[b] | KP22 | WIV[b] |
| CRAB13 | WIV[b] | KP23 | WIV[b] |
| CRAB36 | WIV[b] | KP24 | WIV[b] |
| CRAB27 | WIV[b] | KP26 | WIV[b] |
| CRAB28 | WIV[b] | KP27 | WIV[b] |
| CRAB22 | WIV[b] | KP28 | WIV[b] |
| CRAB26 | WIV[b] | KP602 | WIV[b] |
| CRAB140 | WIV[b] | KP605 | WIV[b] |
| CRAB143 | WIV[b] | KP607 | WIV[b] |
| CRAB147 | WIV[b] | ***Pseudomonas aeruginosa*** | |
| ***Escherichia coli*** | | PAE2555 | WIV[b] |
| *E. coli* 100 | WIV[b] | PAE1 | WIV[b] |
| *E. coli BL21* | WIV[b] | PAE2 | WIV[b] |
| E. coli O91 | WIV[b] | PAE52 | WIV[b] |
| *E. coli* O97 | WIV[b] | PAE53 | WIV[b] |
| *E. coli* O149 | WIV[b] | PAE2139 | WIV[b] |
| *E. coli* O517 | WIV[b] | PAE2169 | WIV[b] |
| *E. coli* MS31 | WIV[b] | PAE2176 | WIV[b] |
| *E. coli* MS33 | WIV[b] | PAE2703 | WIV[b] |
| *E. coli* MS28 | WIV[b] | PAE3223 | WIV[b] |
| *E. coli* MS34 | WIV[b] | PA2-3121 | WIV[b] |
| ***Staphylococcus aureus*** | | PA2-3283 | WIV[b] |
| *S. aureus* N315 | WIV[b] | PA2-3763 | WIV[b] |
| *S. aureus* RN4220 | WIV[b] | PA2-3965 | WIV[b] |
| *S. aureus* NF71 | WIV[b] | ***Streptococcus*** | |
| *S. aureus* 25923 | WIV[b] | *S. pneumoniae* 17NS26 | HPPH[a] |
| *Enterococcus faecalis* | | *S. agalactiae* 12 | WIV[b] |
| *E. faecalis* 493 | WIV[b] | *S. agalactiae* C001 | WIV[b] |
| *E. faecalis* 29212 | ATCC[c] | *S. dysgalactiae* 35666 | ATCC[c] |
| *E. faecalis* 51299 | WIV[b] | *S. pyogenes* 12344 | ATCC[c] |
| *E. faecalis* 571-2 | WIV[b] | *S. mutans* 159 | WIV[b] |
| | | *S. Suis* 1 | HUAHE[d] |

a: Isolated from Henan Provincial People's Hospital, Zhengzhou, China.

b: Obtained from Wuhan Institute of Virology, Chinese Academy of Sciences, Wuhan, China.

c: Purchased from Guangdong Culture Collection Center, China.

d: Obtained from Henan University of Animal Husbandry and Economy, Zhengzhou, China.

**Table S7. Primers used in this study.**

| Primer | Sequence (5'-3') |
|---|---|
| P10-Lys-F | ctttaagaaggagatataccATGggcatgagcaaaaccaccagt |
| P10-Lys-R | gtggtggtgctcgagCAGtttttcagcaggg |
| P11-N-F | agaaggagatataccATGggcatgaccaccaaaccgtttttgatgcag |
| P11-N-R | gtggtggtgctcgagCAGcagtttcagcagggtgcta |
| P11-C-F1 | aactttaagaaggagatataccatgggtggtaaaaccacc |
| P11-C-R1 | ggtggtggtgctcgagtttttcagaaacagtgcacg |
| P11-C-F2 | aactcgagcaccaccaccaccaccac |
| P11-C-R2 | ggtatatctccttcttaaagttaaacaaaattatttctagag |
| K133AR137I-F1 | tttgtttaactttaagaaggagatataccatgggcatgagcaaaaccacc |
| K133AR137I-R1 | acgacggataaccagaccggccataactttaccacc |
| K133AR137I-F2 | tggccggtctggttatccgtcgtgaagcagaac |
| K133AR137I-R2 | gctcatgcccatggtatatctccttcttaaagttaaacaaaattatttc |
| R20IG27D-F1 | accaccagtaatgcaggtctgaatctgatcaaaggttttgaaggtaaaatcctgaatgc |
| R20IG27D-R1 | gtgccaaaaccaatggtccaaacaccaacatcatcatcataggcattcaggattttacc |
| R20IG27D-F2 | ccattggttttggcaccatcaaatatccgaatggtgttcg |
| R20IG27D-R2 | cctgcattactggtggttttgctcatgcccatggtatatc |
| G128DG129D-F1 | atgagcaaaaccaccagtaatgcaggtctgaatctgatcaaag |
| G128DG129D-R1 | aactttatcatcggctttattccaaaccagaaactgatctgctgc |
| G128DG129D-F2 | aagccgatgataaagttatgaaaggtctggttcgtcgtcg |
| G128DG129D-R2 | actggtggttttgctcatgcccatggtatatctccttc |
| G27DR137I-F1 | tgatgatgttggtgtttggaccattggttttggcacc |
| G27DR137I-R1 | tctgcttcacgacggataaccagacctttcataactttaccac |
| G27DR137I-F2 | ggttatccgtcgtgaagcagaacgtgcactg |
| G27DR137I-R2 | ccaaacaccaacatcatcatcataggcattcaggcg |

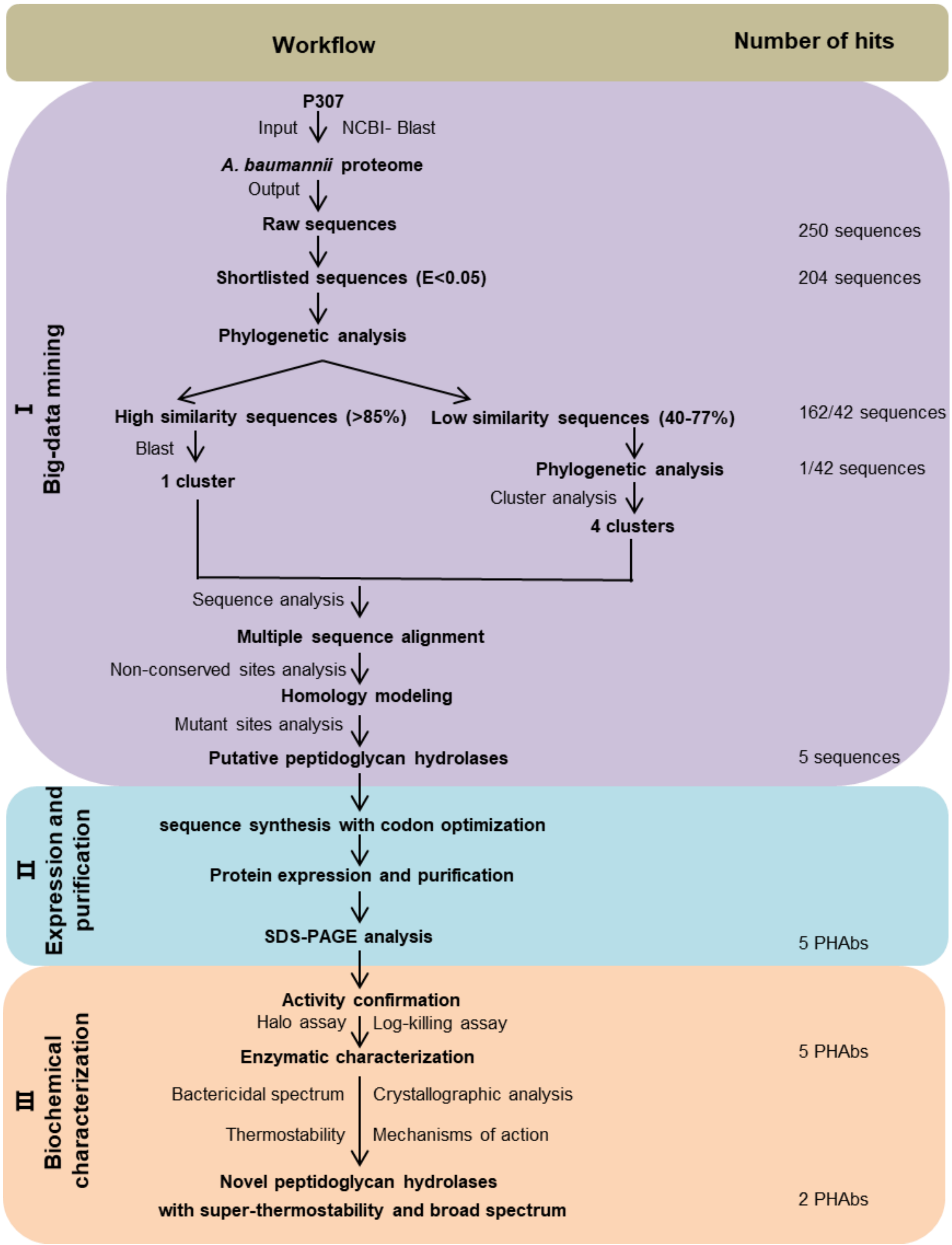


**Figure S1. Workflow of the search integrated system for peptidoglycan hydrolases.** The three different stages of the workflow are highlighted in different colors: (I) Screening of peptidoglycan hydrolases using the cationic peptides P307 as a template (yellow). (II) Protein expression and purification (blue). (III) Biochemical characterization (green).

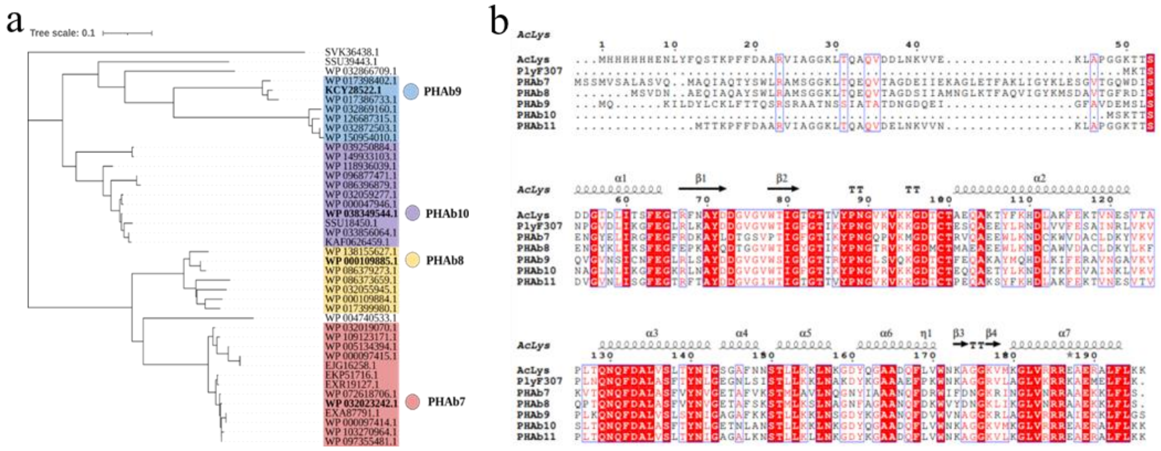


**Figure S2. Selective five putative peptidoglycan hydrolases.** (a) Phylogenetic tree of *A. baumannii* peptidoglycan hydrolases homolog to P307. (b) Multiple sequence alignment of PHAb7, PHAb8, PHAb9, PHAb10, PHAb11, PlyF307, and AcLys using MEGAX software and ClustalW algorithm. Secondary structures are annotated by Espript 3.0 website.

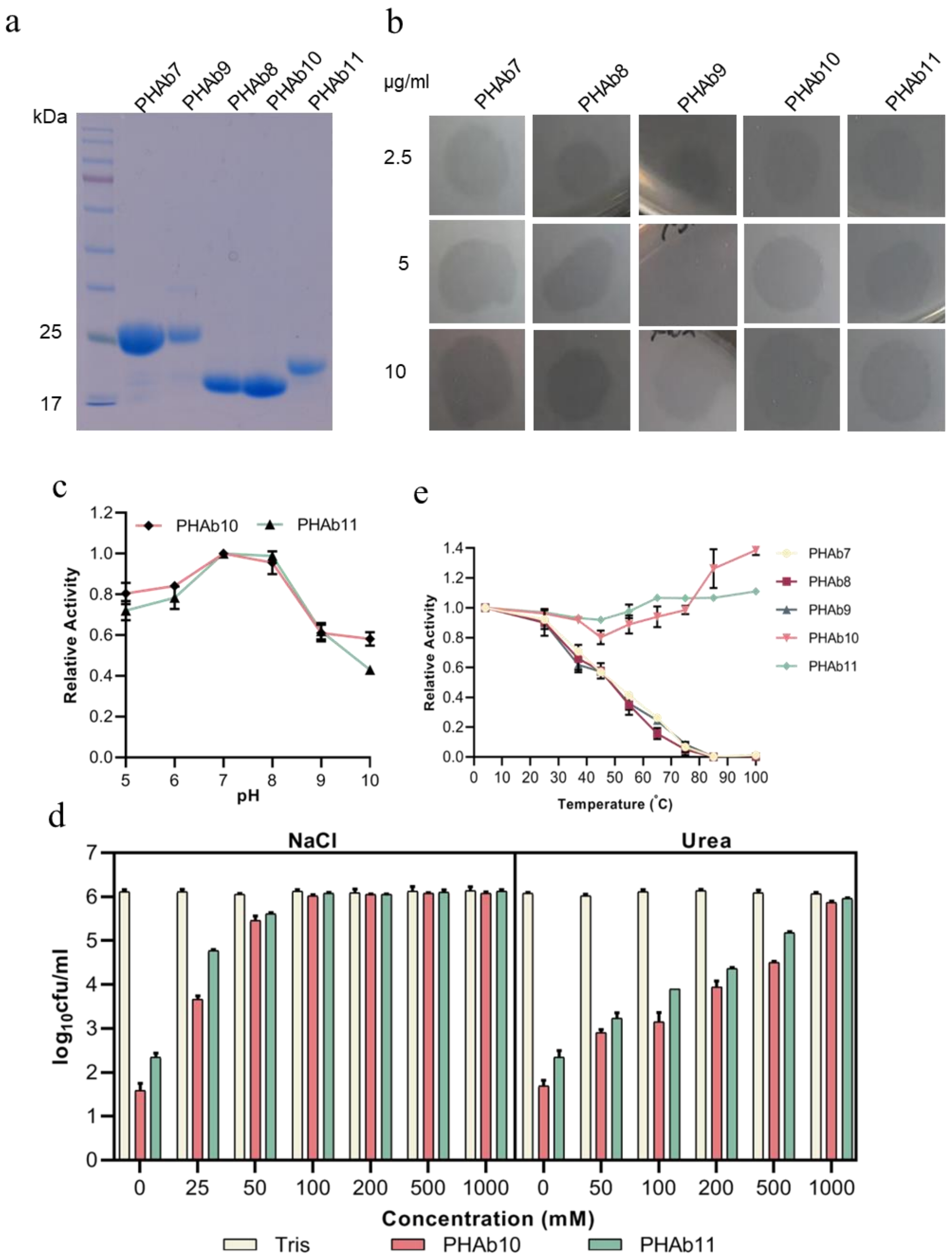


**Figure S3. Testing of the bactericidal activity of the screened putative peptidoglycan hydrolases.** (a) Purified proteins were analyzed on a 12% SDS-PAGE gel. M: standard protein marker. (b) Halo of the five PHAbs on a lawn of *A. baumannii* 3437. The concentrations of each peptidoglycan hydrolase are 2.5 μg/ml, 5 μg/ml, and 10 μg/ml. (c) pH stability of PHAb10 and PHAb11. Log-phase *A. baumannii* 3437 cells were incubated with 50 μg/ml peptidoglycan hydrolases at 37°C for 1 hour at the indicated pH conditions. pH buffer: 20 mM boric acid and 20 mM phosphoric acid were mixed in equal volumes and adjusted to different pH values with HCl. (d) Log-phase *A. baumannii* 3437 cells were incubated with 50 μg/ml peptidoglycan hydrolase in 20 mM Tris-HCl (pH 7.4) in the presence of different concentrations of NaCl (left) and urea (right) for 1 hour at 37°C. (e) Relative activity of five PHAbs against *A. baumannii* 3437 after 1 hour storage at different temperatures. In all cases, viable bacteria were counted by serial dilution and plating assay. Experiments were carried out in triplicate; error bars indicate standard deviation.

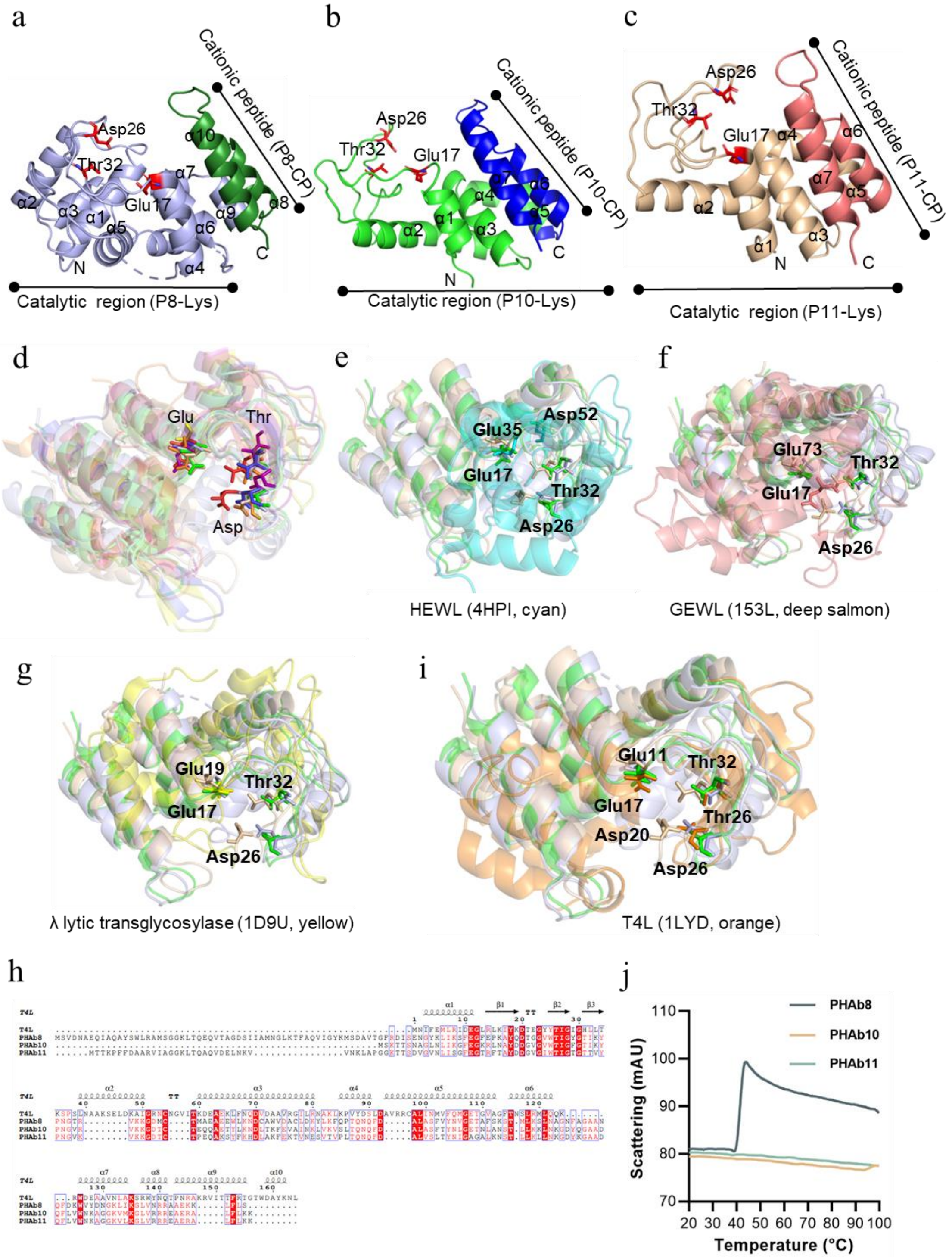


**Figure S4. Structural analysis of PHAb8, PHAb10, and PHAb11.** (a-c) Structures of monomeric PHAb8 (a), PHAb10 (b), and PHAb11 (c). (d) Structural superposition of PHAb8, PHAb10, PHAb11, and five classical T4L-like lysozymes that share high similarity from Dali search. PHAb8 is shown in light blue, PHAb10 in green, PHAb11 in wheat, AcLys in red, LysF1 in blue, SpmX-Mur-*Ae* in yellow, P22 in purple, and R21 in orange. (e-h) Structural alignment of PHAb8, PHAb10, PHAb11 with various lysozymes including HEWL (e, cyan), GEWL (f, deep salmon), phage lambda lytic transglycosylase (g, yellow) and T4 lysozyme (h, orange). Residues in the conserved catalytic triad are shown as sticks. (j) NanoDSF scattering plots of PHAb8, PHAb10, and PHAb11 at different temperatures.

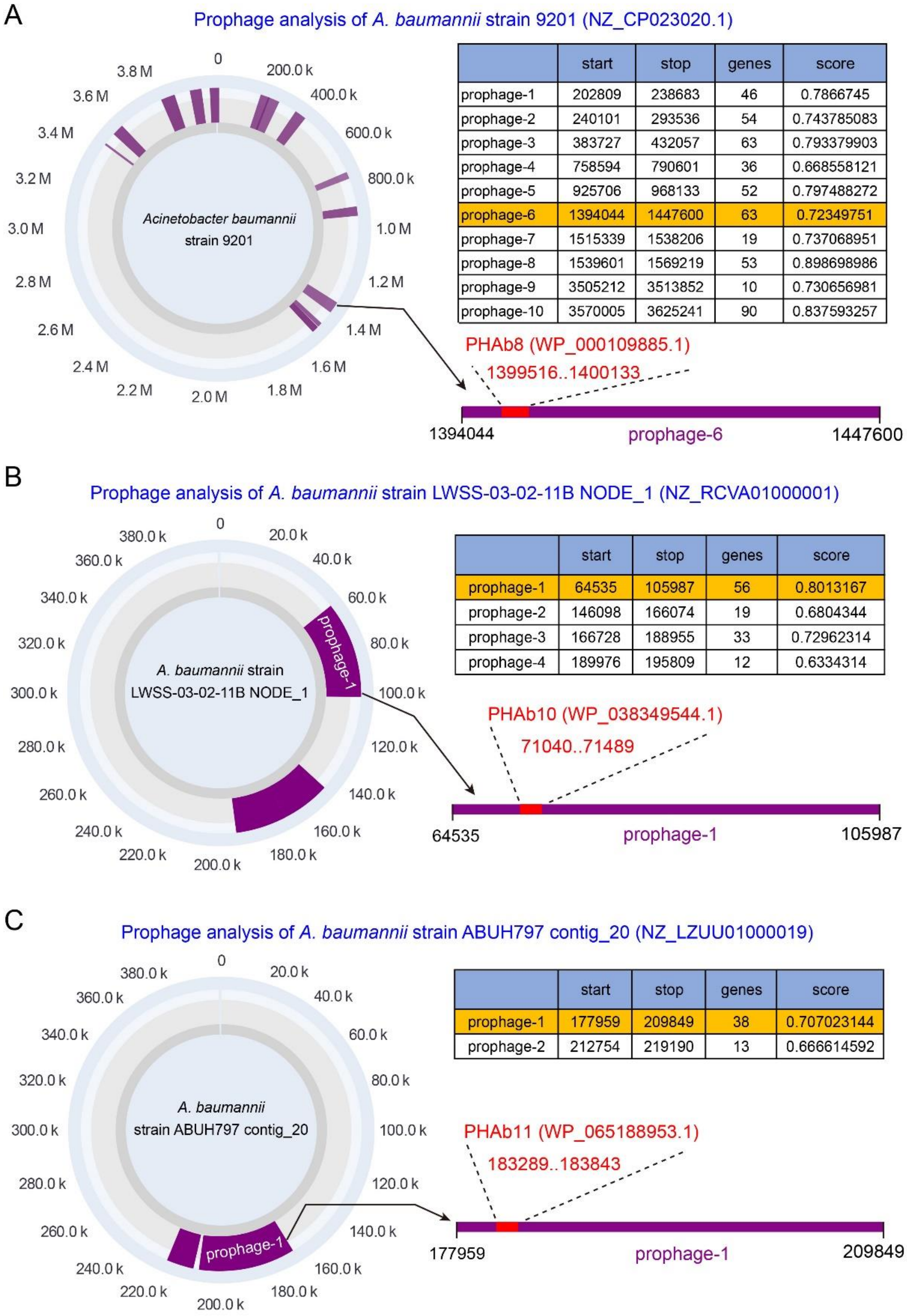


**Figure S5. Prophage analysis of PHAbs.** PHAb8, PHAb10, and PHAb11 encoding sequences were retrieved from *A. baumannii* strain 9201, *A. baumannii* strain LWSS-03-02-11B and *A. baumannii* strain ABUH797 were analyzed by PhageBoost tool. The results show that all three genes are indeed located in the prophage region.

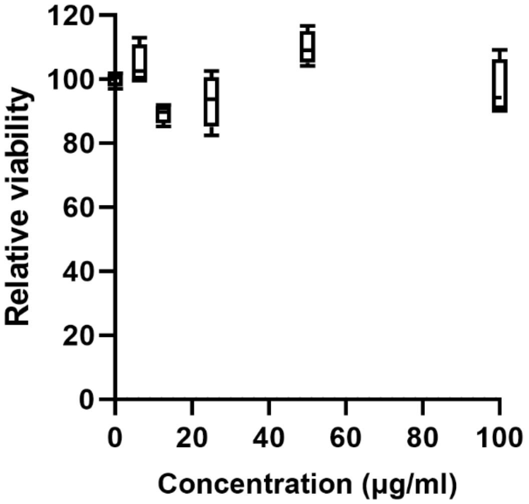


**Figure S6. Relative viability of HepG2 cells exposed to different concentrations of PHAb10.** Cells were cultured in DMEM containing 10% fetal bovine serum, 1% penicillin, and 1% streptomycin for 24 h. After cells were exposed to a series of concentrations of PHAb10 (0, 6.25, 12.5, 25, 50, and 100 μg/ml) for an additional 24 h, the relative viability of cells after each treatment was determined by CCK-8 assay.

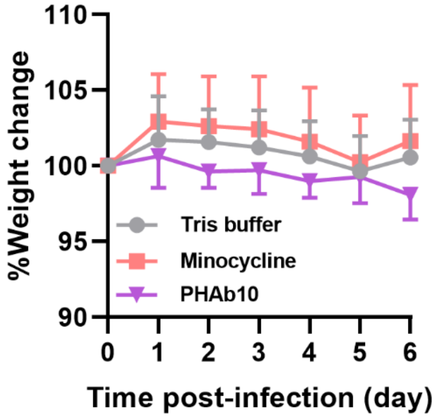


**Figure S7. Changes in mouse body weight in a mouse abscess model.** Mice were subcutaneously injected with 25 µl of $5\times10^8$ cfu/ml of *A. baumannii* 3437 on the right side of the back dorsum. Twenty-four hours after infection, mice were injected subcutaneously with 10 µg/mouse PHAb10 (10 µl; n = 6), 4 µg/mouse minocycline (10 µl; n = 7), or an equal volume of Tris buffer (10 µl; n = 8). Each group was subcutaneously injected once a day for 5 consecutive days. The body weight of mice in each group was monitored daily.
